# PREDICTD: PaRallel Epigenomics Data Imputation with Cloud-based Tensor Decomposition

**DOI:** 10.1101/123927

**Authors:** Timothy J. Durham, Maxwell W. Libbrecht, J. Jeffry Howbert, Jeff Bilmes, William Stafford Noble

## Abstract

The Encyclopedia of DNA Elements (ENCODE) and the Roadmap Epigenomics Project have produced thousands of data sets mapping the epigenome in hundreds of cell types. However, the number of cell types remains too great to comprehensively map given current time and financial constraints. We present a method, PaRallel Epigenomics Data Imputation with Cloud-based Tensor Decomposition (PREDICTD), to address this issue by computationally imputing missing experiments in collections of epigenomics experiments. PREDICTD leverages an intuitive and natural model called “tensor decomposition” to impute many experiments simultaneously. Compared with the current state-of-the-art method, ChromImpute, PREDICTD produces lower overall mean squared error, and combining methods yields further improvement. We show that PREDICTD data can be used to investigate enhancer biology at non-coding human accelerated regions. PREDICTD provides reference imputed data sets and open-source software for investigating new cell types, and demonstrates the utility of tensor decomposition and cloud computing, two technologies increasingly applicable in bioinformatics.

## 1 Introduction

Understanding how the genome is interpreted by varied cell types in different developmental and environmental contexts is a key question in biology. With the advent of high throughput next generation sequencing technologies, over the past decade we have witnessed an explosion in the number of assays to characterize the epigenome and interrogate chromatin state genome-wide. Assays to measure chromatin accessibility (DNase-seq, ATAC-seq, FAIRE-seq), DNA methylation (RRBS, WGBS), and histone modification and transcription factor binding (ChIP-seq) have been leveraged in large projects such as the Encyclopedia of DNA Elements (ENCODE) [1] and the Roadmap Epigenomics Project [2] to characterize patterns of biochemical activity across the genome in many different cell types and developmental stages. These projects have produced thousands of genome-wide data sets, and studies leveraging these data sets have provided insight into multiple aspects of genome regulation, including mapping different classes of genomic elements [3, 4], inferring gene regulatory networks [5], and providing insights into possible disease-causing mutations identified in genome-wide association studies [2].

Despite the progress made by these efforts to map the epigenome, much work remains to be done. Due to time and funding constraints, data have been collected for only a fraction of the possible pairs of cell types and assays defined in these projects (Fig. 1A). Furthermore, taking into account all possible developmental stages and environmental conditions, the number of possible human cell types is nearly infinite, and it is clear that we will never be able to collect data for all cell type/assay pairs. However, understanding the epigenome is not an intractable problem because in reality many of the assays detect overlapping signals such that most of the unique information can be recovered from just a subset of experiments. One solution is thus to prioritize experiments for new cell types based on analysis of existing data [6]. Alternatively, one may exploit existing data to accurately impute the results of missing experiments.

Ernst and Kellis pioneered this imputation approach, and achieved remarkable accuracy with their method, ChromImpute [7]. Briefly, this method imputes data for a particular target assay in a particular target cell type by 1) finding the top ten cell types most correlated with the target cell type based on data from non-target assays, 2) extracting features from the data for the target assay from the top ten non-target cell types, and also extracting features from the data for nontarget assays in the target cell type, and 3) training a regression tree for each of the top ten most correlated cell types. Data points along the genome are imputed as the mean predicted value from the collection of trained regression trees. Although ChromImpute produces highly accurate imputed data, this training scheme is complicated, and results in a fragmented model of the epigenome that is very difficult to interpret. We hypothesized that an alternative approach, in which a single joint model learns to impute all experiments at once, would simplify model training and improve interpretability while maintaining accurate imputation of missing data.

Accordingly, we present PaRallel Epigenomics Data Imputation using Cloud-based Tensor Decomposition (PREDICTD), which treats the imputation problem as a tensor completion task and employs a parallelized algorithm based on the PARAFAC/CANDECOMP method [8, 9]. Our implementation, which runs on consumer cloud infrastructure, achieves high-accuracy imputation of ENCODE and Roadmap Epigenomics data and predicts all data sets jointly in a single model. We explain the model, discuss its performance on held out experiments from the Roadmap Epigenomics Consolidated data [2], show that the model parameters summarize biologically relevant features in the data, and demonstrate that imputed data can recapitulate important cell type-specific gene regulatory signals in non-coding human accelerated regions of the genome [10].

## 2 Results

### 2.1 Epigenomic maps can be imputed using tensor factorization

Data from the Roadmap and ENCODE projects can be organized into a three-dimensional tensor, with axes corresponding to cell types, assays, and genomic positions (Fig. 1B). This tensor is long and skinny, with many fewer cell types and assays than genomic positions, and the data for experiments that have not been done yet are missing in the tensor fibers along the genome dimension. Our strategy for imputing these fibers is to jointly learn three factor matrices that can be combined mathematically to produce a complete tensor that both approximates the observed data and predicts the missing data. These three factor matrices are of shape C × *L, A* × *L*, and *G × L*, where *C, A*, and *G* indicate the numbers of cell types, assays, and genomic positions, respectively, and *L* indicates the number of “latent factors” that the model trains (Fig. 1B), and thus the number of model parameters.

We developed and trained our implementation of this tensor factorization model, PREDICTD, using 1014 data sets from the Roadmap Epigenomics Consolidated Epigenomes [2] (Fig. 1A). To assess model performance, we split the data sets into five training/test splits, and we report on the results of imputing each test set at 25 base pair resolution. The model training proceeds by distributing the data and genome parameters across the nodes of the cluster, and then sharing the cell type and assay parameters across all nodes using a parallelized training procedure (See Methods, Fig. 1C). We find that training on a randomly selected 0.01% of the genome provides enough signal for learning the cell type and assay parameters (Fig. S2); these parameters are then applied across all genomic positions of interest by training the genome parameters for each position while holding the cell type and assay parameters constant. We report results from imputing just over 1% of the genome, including the ENCODE Pilot Regions [1] and 2,640 non-coding human accelerated regions [10], and all references to the genome dimension refer to this subset of loci.

Our model formulation and implementation offer several important advantages. First, training a single model to impute all data sets at once is a straightforward and intuitive way of solving this problem. Second, as we demonstrate below, the model can leverage the joint training to perform well even on cell types with a single informative experiment. Third, the parameters of the trained model have the same semantics across all input data sets and can be interrogated to learn about relationships among assays, cell types, and genomic loci. Last, PREDICTD software is open source (https://github.com/tdurham86/PREDICTD.git), and is also implemented and distributed on the consumer cloud, which makes our model immediately accessible to and easily runnable by nearly anyone.

### 2.2 PREDICTD imputes epigenomics experiments with high accuracy

PREDICTD imputes missing data with high accuracy based on both visual inspection and quality measures (Fig. 2). Visually, the imputed signal pattern closely matches that of observed data, and recapitulates the known associations of epigenomic marks with genomic features (Fig. 2A, S1). For example, as expected, H3K4me3 imputed signal is strongly enriched in narrow peaks at promoter regions near the transcription start site of active genes, and H3K36me3, known to mark transcribing regions, is enriched over gene bodies.

We also show strong performance of PREDICTD on ten different quality measures (see Methods, Figs. S3, S8), especially the global mean squared error metric (MSEglobal). As a key part of the PREDICTD objective function, MSEglobal is explicitly optimized during model training. The MSEglobal measure has a mean of 0.1251, and it ranges from 0.0375 for H3K9me3 in the “CD3 Primary Cells Cord Blood” cell type to 0.4437 for H4K20me1 in “Monocytes-CD14+ RO01746”. Other key quality measures include the genome-wide Pearson correlation (GWcorr, mean: 0.6825, min: 0.0623 for H3K9me3 in “ES-WA7 Cell Line”, max: 0.9337 for H3K4me3 in “HUES64 Cell Line”), and the area under the receiver operating characteristic curve for recovering observed peak regions from imputed data (CatchPeakObs, mean: 0.9558, min: 0.5683 for H3K36me3 in “Right Atrium”, max: 0.9982 for H3K4me3 in “NHLF Lung Fibroblasts”). Note that seven of our ten quality measures, including GWcorr and CatchPeakObs, were also used in the ChromImpute publication [7].

As a baseline, we compared the performance of PREDICTD to a simple “main effects” model, which computes the global mean of the observed data, and then the column and row residuals of each two-dimensional slice of the tensor along the genome dimension, and imputes a given cell in the tensor by summing the global mean and the corresponding row and column residual means. PREDICTD outperforms this baseline model for MSEglobal on all but three assays (Fig. 2B).

Furthermore, PREDICTD similarly outperforms the main effects on all additional performance metrics (Fig. S3A).

**Figure 1:**
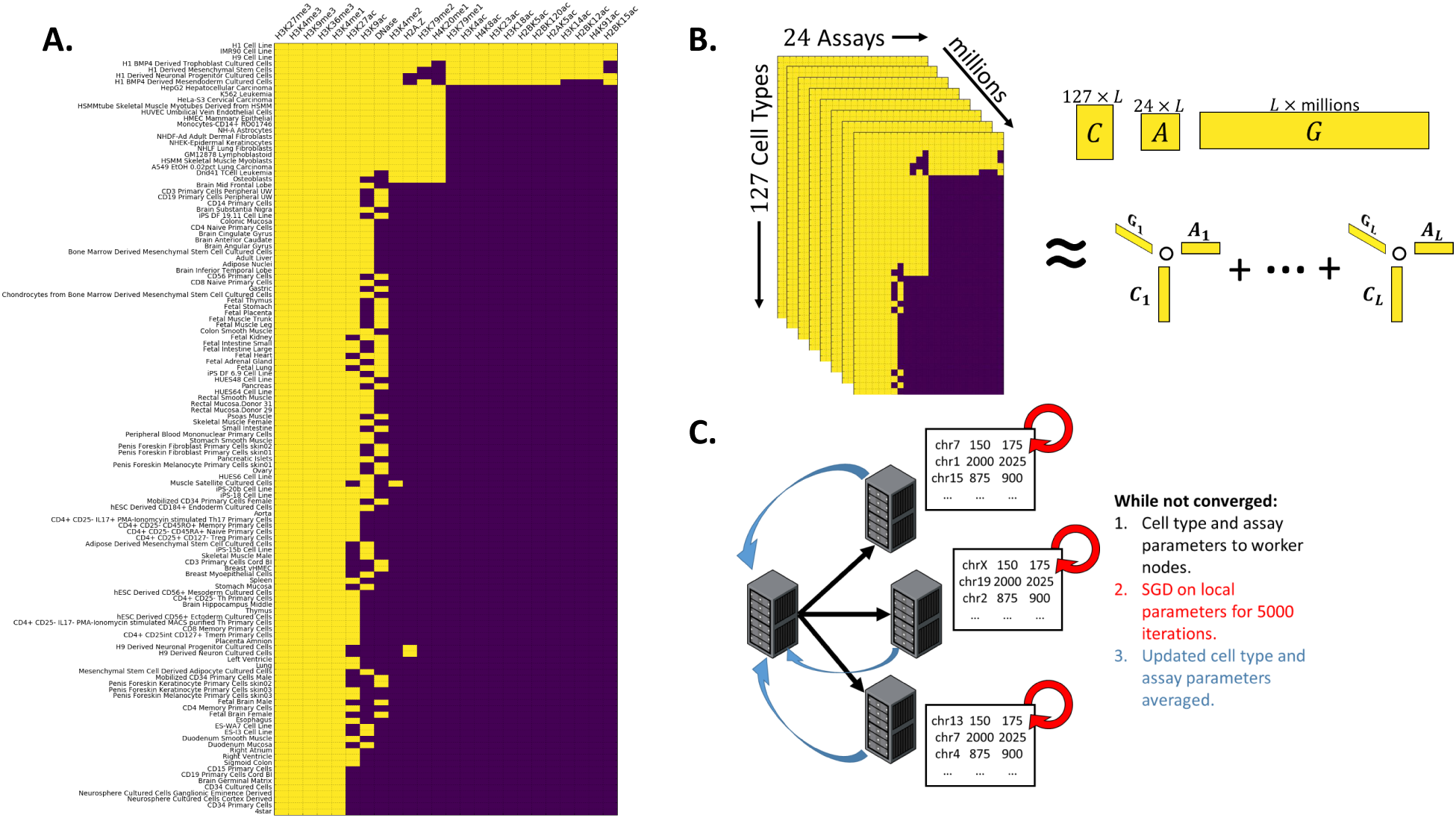
Overview. **A.** Matrix representing the subset of the Roadmap Epigenomics consolidated data set used in this study. Experiments in yellow have observed data, while missing experiments are purple. **B.** We model the experiments in A. as a three-dimensional tensor, and find three low-rank factor matrices (denoted *C, A*, and *G*) that can be combined by summing the outer products of each of the *L* latent factor vector triplets to reconstruct a complete tensor with no missing values that both approximates the existing data and imputes the missing data. **C.** The genome dimension is very large, so in order to fit all of the data in memory and to speed up training, we distribute the tensor across multiple cluster nodes running Apache Spark. Then we use parallel stochastic gradient descent [11] to share the *A* and *C* matrices across all nodes.

**Figure 2:**
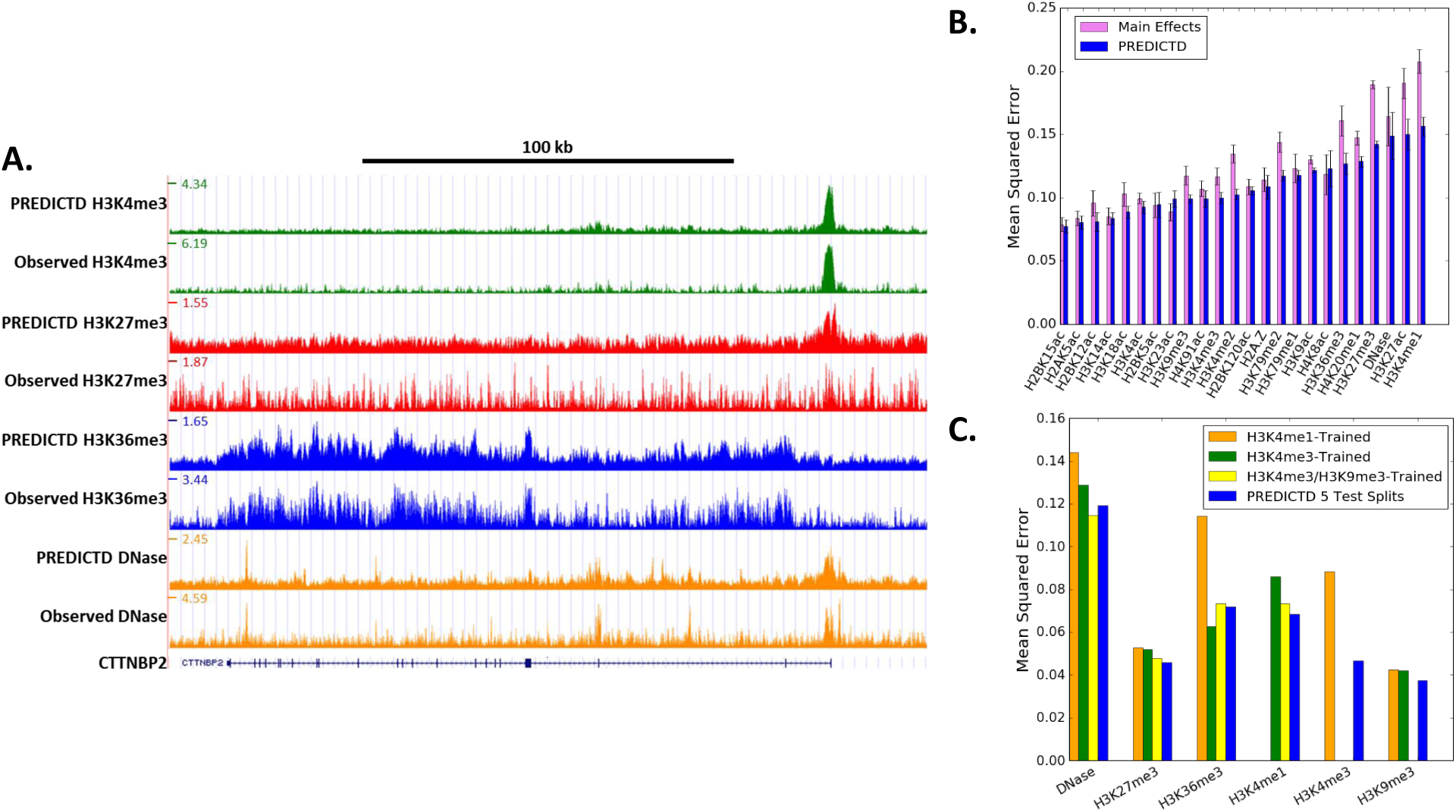
PREDICTD imputes missing epigenomics data with high accuracy. **A.** Comparison of PREDICTD and observed data in H1 embryonic stem cells, which is one of three cell types with observed data for all assays. For each assay, the top track is PREDICTD and the bottom is observed. See Fig. S1 for more assays**. B.** Global mean squared error (MSEglobal) for each data set, averaged by assay for the baseline main effects model (pink), and PREDICTD (blue). PREDICTD outperforms the main effects on all but three assays. **C.** Simulated analysis of a cell type with few available data sets. The bar graph shows the MSEglobal measure for the six available assays for the “CD3 Primary Cells from Cord Blood” cell type on four models. In blue is the full PREDICTD model shown in A and B, while the other bars represent models trained with all observed data, but only H3K4me1 (orange), only H3K4me3 (green), or only H3K4me3 and H3K9me3 (yellow) from “CD3 Primary Cells from Cord Blood”.

### 2.3 PREDICTD can accurately impute data for under-characterized cell types

A key application of PREDICTD will be to impute results for cell types that may have only one or two data sets available. To investigate the performance of PREDICTD in this context, we trained a model on all available data for all cell types, except that we only included one or two experiments for the “CD3 Cord Blood Primary Cells” cell type. In particular, one model had just H3K4me1 in the training set for this cell type, one had just H3K4me3, and one had both H3K4me3 and H3K9me3. Comparing the performance metrics between these experiments and the imputed results from our original models trained on the five test folds, we find that the results of training with just H3K4me3 or both H3K4me3 and H3K9me3 are nearly as good as (and sometimes better than) the results from the original models with training data including five or six experiments for this cell type (Fig. 2C). Imputing only based on H3K4me1 signal did not perform as well as imputing based on only H3K4me3. This observation is consistent with previous results on assay prioritization [6] indicating that H3K4me3 is the most information-rich assay. We conclude that PREDICTD performs well on under-characterized cell types and will be useful for studying new cell types for which few data sets are present.

### 2.4 Model parameters capture patterns that summarize information about cell types, assays, and genomic loci

Although it is not straightforward to assign semantics to individual latent factors, we find that analyzing their values in aggregate shows patterns that recapitulate known relationships among the cell types and assays and that associate with particular types of genomic elements (Fig. 3). Hierarchical clustering on the rows of the cell type factor matrix shows that similar cell types are grouped together, producing large clades for cell lines, embryonic stem cells, immune cell types, and brain tissues (Fig. 3A, S4). In the same way, clustering groups together assays with similar targets, for example, a clade for more punctate promoter marks and another for broader marks that are found at transcribing genes or heterochromatin (Fig. 3B). Furthermore, these clustering results are highly non-random. We quantified this by comparing our clustering results to randomly shuffled cluster identities using the Calinski-Harabaz Index, which assesses how well the clustering separates the data by comparing the average distance among points between clusters to the average distance among points within clusters (Fig. S5).

**Figure 3:**
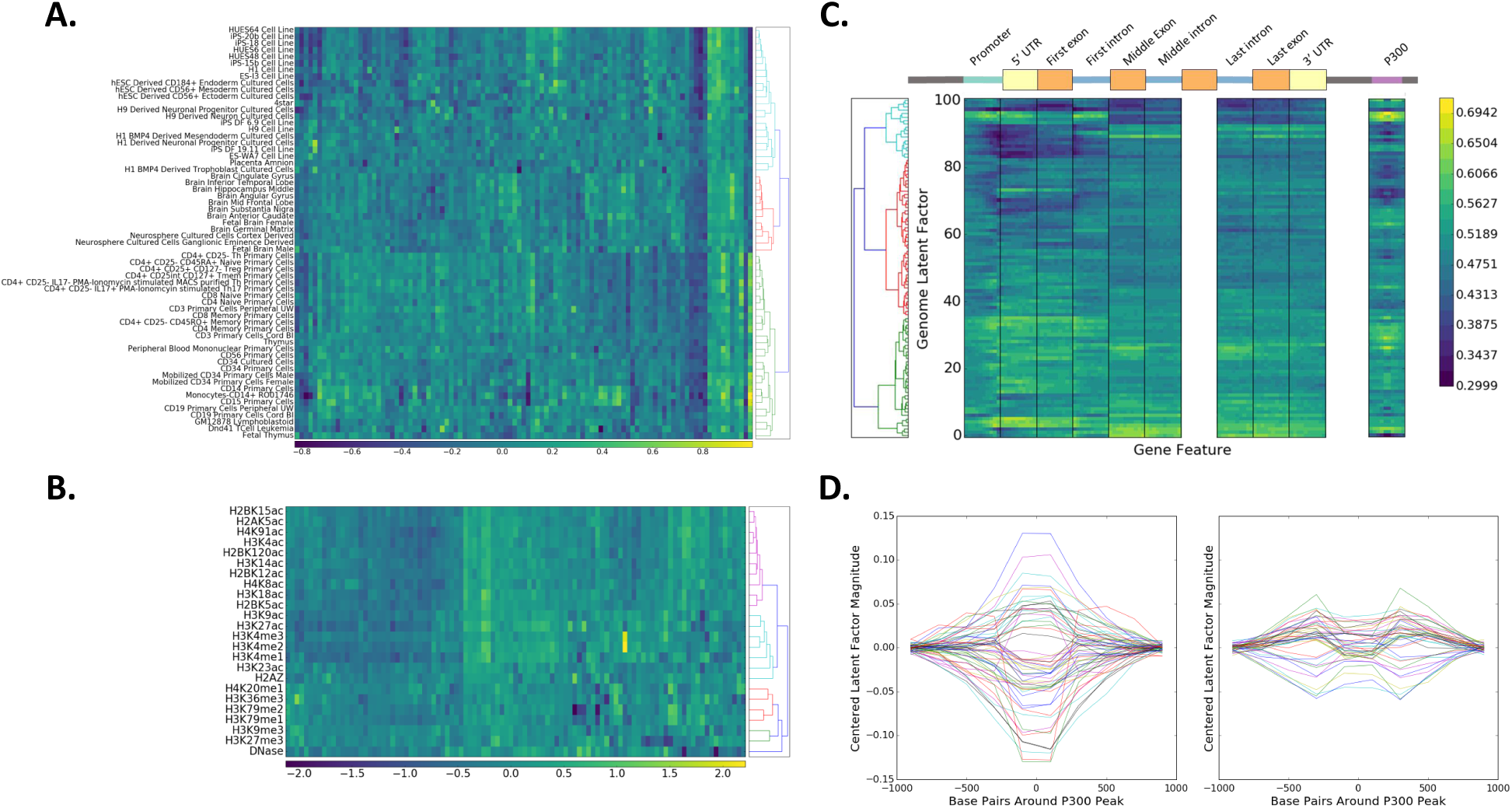
The model parameters capture meaningful relationships relative to each tensor dimension. **A.** Heatmap showing cell type latent factor values from one of the trained models. Hierarchical clustering of the cell types shows that similar cell types tend to cluster together, with particularly prominent clusters for brain (red), immune (green), and embryonic stem cell types (cyan). This is a subset of cell types; for the full clustering see Fig. S4**. B.** Heatmap showing assay latent factor values from one of the trained models. Hierarchical clustering shows that similar assays tend to cluster together, with the colored clades from top to bottom representing acetylation marks generally associated with active promoters (purple), marks strongly associated with active regulatory regions (cyan), broad marks for transcription (red) and repression (green), and finally DNase hypersensitivity**. C.** Latent factor values show patterns particular to different parts of the gene and to P300 peaks called from ENCODE data**. D.** Average parameter values for each latent factor plotted as a lines spanning the region +/- 1kb around the center of P300 peaks. Parameter values are centered at zero and plotted based on whether they show the highest magnitude at the peak (left, 59 latent factors) or flanking the peak (right, 42 latent factors).

For the genome factor matrix, we projected the coordinates of each gene from the GENCODE v19 human genome annotation [12] (https://www.gencodegenes.org/releases/19.html) onto an idealized gene model that includes nine parts from 5’ to 3’ in the gene: the promoter, 5’ untranslated region (UTR), first exon, first intron, middle exons, middle introns, last intron, last exon, and 3’ UTR. This procedure produced a summary of the genome latent factors (Fig. 3C) that clearly shows distinct patterns among the latent factors at different gene components. For example, some latent factors tend to have high values at regions near the transcription start site and 5’ end of the gene, while others have low values. We also find particular patterns in the latent factors that are associated with exons and introns.

In addition to investigating patterns in the genome latent factors at genes, we checked for a signature of distal regulatory regions. P300 is a chromatin regulator that associates with active enhancers [13]. We therefore decided to search for patterns in the genome latent factors at windows +/- 1 kb around annotated P300 sites from published ENCODE tracks (see Methods). Note that no P300 data was used to train PREDICTD. Nonetheless, we find a striking pattern, with clear signal within the 400 bp region surrounding the center of the peak, and some latent factors even show a flanking pattern in the bins 200-400 bp away from the peak center (Fig. 3C,D). If we randomize the latent factors at each genomic location and do these same analyses we find no discernible pattern (Fig. S6). We thus conclude that the trained model parameters encode patterns that correspond to biology.

### 2.5 PREDICTD and ChromImpute results are similar and complementary

As described above, the ChromImpute method [7] provides high quality imputed data but employs a complicated model and training procedure tuned to each individual experiment. In contrast, our tensor decomposition approach imputes all missing experiments by using a single model, which we argue is conceptually simpler and addresses the problem in a more natural way. Furthermore, we find that our model outperforms ChromImpute on our primary performance measure (MSEglobal), and yields similar performance on nine additional measures (Fig. 4A, Table 1, and Fig. S8). The correlation of quality measures between PREDICTD and ChromImpute is higher than the correlation between the main effects method and ChromImpute, indicating that PREDICTD agrees with ChromImpute more often than main effects does. Furthermore, the mean log ratio of quality measures on corresponding experiments imputed by PREDICTD and ChromImpute show smaller differences than the log ratios for main effects and ChromImpute (Table 1, Fig. 4). Thus, PREDICTD produces high quality imputed data that is almost as good or better than ChromImpute predictions, depending upon which quality measure is employed.

**Figure 4:**
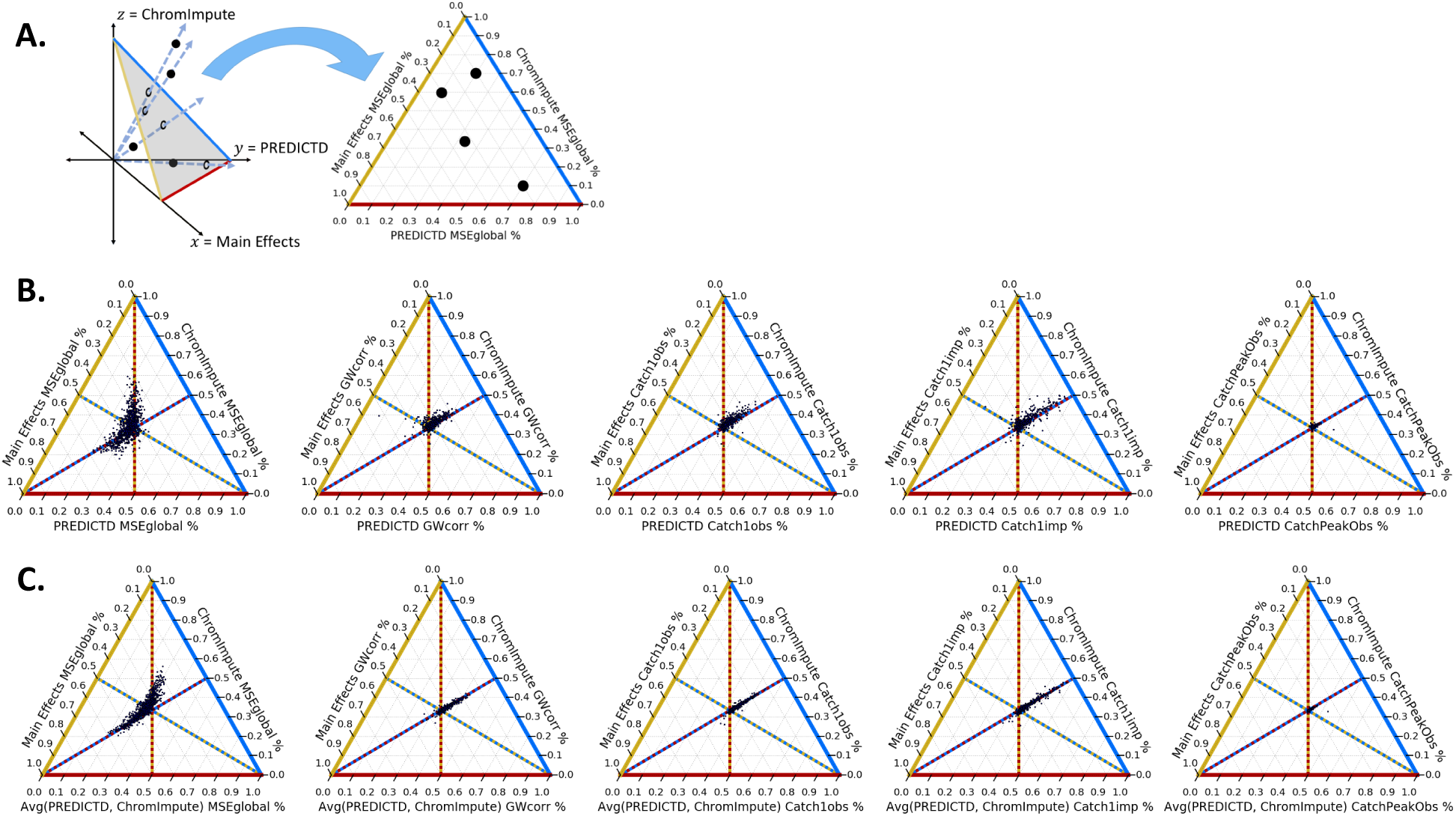
PREDICTD performs comparably to ChromImpute, and combining the models improves the result. A. Schematic describing how a ternary plot relates to Cartesian coordinates. B. Comparing PREDICTD, ChromImpute, and main effects models across five quality measures: the global mean squared error (MSEglobal), the genome-wide Pearson correlation (GW-corr), the percent of the top 1% of observed data windows by signal value found in the top 5% of imputed windows (Catch1obs), the percent of the top 1% of imputed windows by signal value found in the top 5% of observed windows (Catch1imp), and the area under the receiver operating characteristic curve for recovery of observed peak calls from all imputed windows ranked by signal value (CatchPeakObs). Dots at the center of the plot indicate experiments for which the three models perform comparably, while a spread of points out from the center indicates that the models perform differently for those experiments. Lines bisecting the triangles indicate axes over which the performance of two models can be compared, and which two models to compare is indicated by the colors of each line. For example, for any level of performance of the main effects model (gold), if the PREDICTD (red) and ChromImpute (blue) models perform comparably, dots will cluster along the red and blue-dotted line. Furthermore, dots representing experiments for which PREDICTD outperforms ChromImpute will fall above this line for MSEglobal (lower is better) and below for the other quality measures (higher is better). Analogous rules apply for comparing the other two pairs of models. C. The same as in A, except that the quality measures for the averaged results of ChromImpute and PREDICTD are plotted along the bottom axis instead of the measures for PREDICTD alone.

We also calculated the distribution of the differences between imputed values and observed values for experiments imputed by both PREDICTD and ChromImpute, and we found that ChromImpute tends to impute higher values than PREDICTD (Fig. S7). We hypothesized that the two models each perform better on different parts of the genome, and so we tried averaging the PREDICTD and ChromImpute results. By the MSEglobal measure, we do see a significant improvement relative to both models, and other metrics on which ChromImpute out-performed PREDICTD alone show parity between ChromImpute and the averaged model. (Fig. 4B).

**Table 1:** Statistics comparing models across five quality measures. See Supplementary Table 1 for the statistics on all quality measures.

| Measure | PREDICTD<br>vs<br>ChromImpute |  |  | PREDICTD<br>vs<br>Main Effects |  |  | Main Effects<br>vs<br>ChromImpute |  |  |
| --- | --- | --- | --- | --- | --- | --- | --- | --- | --- |
|  | corr | log ratio |  | corr | log ratio |  | corr | log ratio |  |
|  |  | mean | std |  | mean | std |  | mean | std |
| **MSEglobal** | 0.691 | -0.131 | 0.265 | 0.841 | -0.168 | 0.209 | 0.510 | 0.037 | 0.373 |
| **GWcorr** | 0.977 | -0.049 | 0.083 | 0.881 | 0.091 | 0.170 | 0.866 | -0.139 | 0.164 |
| **Catch1obs** | 0.977 | -0.035 | 0.056 | 0.921 | 0.091 | 0.156 | 0.886 | -0.125 | 0.177 |
| **Catch1imp** | 0.972 | -0.033 | 0.080 | 0.879 | 0.145 | 0.293 | 0.848 | -0.178 | 0.306 |
| **CatchPeakObs** | 0.922 | -0.008 | 0.017 | 0.779 | 0.024 | 0.037 | 0.812 | -0.032 | 0.035 |

### 2.6 Imputed data can recover cell type-specific enhancer signatures at noncoding human accelerated regions

Human accelerated regions (HARs) are genomic loci that are highly conserved across mammals but harbor more mutations in human than would be expected for their level of conservation (reviewed in [14]). Although some HARs overlap coding regions, the overwhelming majority (>90%) are found in non-coding portions of the genome (non-coding human accelerated regions, or ncHARs) [10, 14], and ncHARs are thought to be enriched for mutations that affect the regulation of genes underlying human-specific traits. Non-coding variation is thought to account for much of our phenotypic divergence from other primates [15], and additional evidence in support of this hypothesis comes from observations that ncHARs cluster around developmental and transcription factor genes [10, 14], transgenic assays for functional validation of enhancer activity [10, 16–18], and computational epigenomics and population genetics studies [10, 19, 20].

In particular, ncHARs are enriched in developmental enhancer activity [10, 19]. In [10], EnhancerFinder [19], a program to predict genomic regions with tissue-specific developmental enhancer activity, was trained on ENCODE epigenomics maps [1] and results from the VISTA enhancer database [21], and applied to ncHARs. EnhancerFinder predicted enhancer activity for 773 of 2649 ncHARs, but the authors note that the characterization of these regions remains incomplete due to limitations in the available data. To our knowledge, no one has yet analyzed enhancer signatures of ncHARs in the context of the Epigenomics Roadmap data. Thus, we decided to address this question as a way to validate PREDICTD in a biological application and to extend the EnhancerFinder results by assessing cell type-specific enhancer activity in the ncHARs based on the Roadmap data set.

Briefly, we imputed data for three enhancer-associated assays (DNase, H3K27ac, and H3K4me1) in all cell types, and averaged the imputed signal over each ncHAR to produce a small tensor with axes corresponding to three assays, 2640 ncHARs, and 127 cell types. We flattened the assay dimension of this tensor by taking the first principal component, then used a biclustering algorithm to group the ncHARs and cell types (see Methods). The resulting cell type groups are consistent with tissue of origin (Fig. 5A, Supplementary Table 2), and the ncHARs cluster based on enhancer-associated signal in different cell type clusters as follows: No signal (65% of the ncHARs), Brain/ES 1 and Brain/ES 2 (25%), Epithelial/Mesenchymal (7%), Non-immune (2%), and Immune (1%) (Fig. 5A, Supplementary Table 3). Using the same strategy to cluster the available observed data gives very similar results, as quantified by the adjusted Rand index (Fig. 5B), especially when compared to two background models: Shuffled, in which the ncHAR coordinates have been randomly shuffled along the genome; and Other, in which the enhancer-associated marks were exchanged for three non-enhancer-associated marks (H3K4me3, H3K27me3, H3K36me3). A heatmap showing the clustering of observed data is provided in Fig. S10.

**Figure 5:**
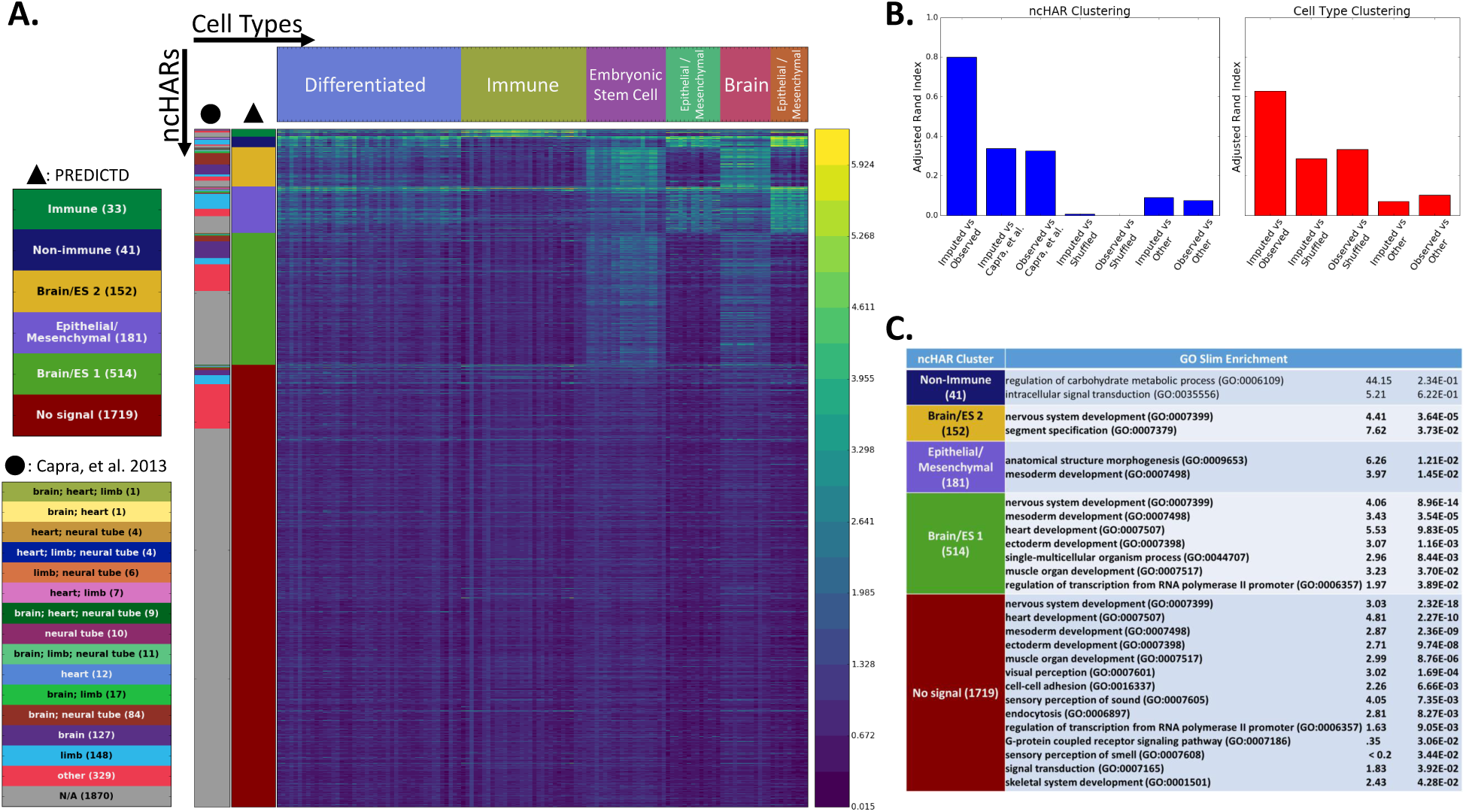
Imputation of enhancer marks reveals tissue-specific patterns of enhancer-associated marks at non-coding human accelerated regions (ncHARs). **A.** Average PREDICTD signal at each ncHAR was compiled for H3K4me1, H3K27ac, and DNase assays from all cell types. The first principal component with respect to the three assays was used in a biclustering to find 6 clusters each along the cell type and ncHAR dimensions. The inverse hyperbolic sine-transformed signal from each of these assays was summed per cell type and ncHAR, and the resulting values were plotted as a heat map. The column marked with a black triangle at the top designates the color key for the ncHAR clusters. The leftmost column, designated with a black circle, identifies ncHARs with predicted tissue-specific developmental enhancer activity based on EnhancerFinder analysis from Capra et al, 2013. **B.** Evaluation of the clustering results with the adjusted Rand index. The clustering results for observed data and PREDICTD for the ncHAR and cell type clusterings, and also those from Capra, et al. 2013 for the ncHARs, all show higher adjusted Rand index scores than the clustering results for observed data with shuffled ncHAR coordinates (Shuffled) or for observed data from non-enhancer-associated marks (Other). **C.** Using the nearest genes for the ncHARS in each cluster, we found that the genes in five of the six clusters deviated from the genomic background based on GO Biological Process overrepresentation analysis. Terms with Bonferroni-corrected p<0.05 are in bold text. Clusters not listed had no GO terms with p-values less than one.

These biclustering results also agree with and expand upon previously published tissue specificity predictions from EnhancerFinder [10, 19]. The brain enhancer predictions from that study are visibly enriched in our Brain/ES clusters, and limb and heart predictions are enriched in our clusters showing activity in differentiated, epithelial, and mesenchymal cell types (Fig. 5A). If we treat the EnhancerFinder tissue assignments [10] as another clustering of the ncHARs, we find that they are more similar to our clustering (both for observed and imputed data) than to either background clustering (Fig. 5B). In addition, our results expand on EnhancerFinder by assigning to cell type-associated clusters 555 ncHARs (21% of ncHARs) characterized by EnhancerFinder as either having activity in “other” tissues (156 ncHARs) or no developmental enhancer activity (“N/A”, 399 ncHARs). We also find that our clustering successfully predicts enhancer activity for many functionally validated ncHARs, and furthermore assigns most of them to the correct cell types (Supplementary Table 4). Briefly, we correctly identify enhancer activity in 14 of 23 ncHARs with evidence in the VISTA database[10, 21], and 6 of 7 ncHARs with validation results suggesting enhancer activity specific to the human allele and not the chimp allele [10]; we find evidence of enhancer identity for three ncHARs associated with *AUTS2*, a gene associated with autism spectrum disorder, including two that showed transgenic enhancer activity [18]; *NPAS3* is a gene associated with schizophrenia that lies in a large cluster of 14 ncHARs, and we find enhancer signal for 8 of 11 of those that have validated enhancer activity [17]; last, HAR2 is a ncHAR with validated human-specific limb enhancer activity that clusters with our Brain/ES category [16]. Thus, assessing potential enhancer activity based on the Roadmap Epigenomics data, which encompasses different cell types and developmental stages than ENCODE, agrees with previous results and expands on them to characterize more ncHARs as having potential tissue-specific enhancer activity.

Finally, we asked what types of biological processes these putative enhancers might regulate. We identified the nearest gene for each ncHAR and conducted a Gene Ontology (GO) biological process term enrichment analysis on our clusters [22–25]. Five of the six clusters showed some level of term enrichment, and four of those five clusters showed terms with enrichment over whole genome background (Bonferonni-corrected *p* < 0.05, Fig. 5C, S11). Note that ncHARs generally tend to cluster near genes involved in development [10], which explains the enrichment of developmental terms even in the case of our No signal cluster. However, we find it encouraging that despite this bias and our relatively small gene lists, the ncHARs that cluster based on tissue-specific enhancer-associated activity show different enriched GO terms that are consistent with their cell type assignments.

Taken together, these results show that PREDICTD imputed data can capture cell type-specific regulatory signals and that PREDICTD can be used as a tool to study the biology of new and under-characterized cell types in the future.

## 3 Discussion

PREDICTD imputes thousands of epigenomics maps in parallel using a three-dimensional tensor factorization model. Our work makes several important contributions. First, the model leverages a machine learning method, tensor decomposition, that holds particular promise in genomics for analyzing increasingly high dimensional data sets. Tensor factorization with the PARAFAC/CANDECOMP procedure was first proposed by two groups independently in 1970 in the context of analyzing psychometric electroencephalogram (EEG) data [8, 9]. Tensor decomposition by this and related methods has since been applied in many other fields [26, 27], and increasingly in biomedical fields as well [28, 29]. Tensor decomposition has advantages over twodimensional methods because taking into account more than two dimensions reduces the rotational flexibility of the model and helps drive the factors to a solution that can explain patterns in all dimensions at once. Our particular application, completing a tensor with missing data, is an area of active research [30] and is analogous to methods for factor matrixization that have proven effective in machine learning applications like recommender systems [31]. To our knowledge, PREDICTD is just the second application of the tensor decomposition approach to epigenomics data [29], and the first to use a tensor completion approach to impute missing data. As such, our method demonstrates another way forward for integrating and jointly analyzing increasingly large and complex data sets in the field.

We compared imputed data from PREDICTD to the published imputed data from ChromImpute [7]. PREDICTD data sets outperform ChromImpute on the global mean squared error metric, but generally slightly under-perform ChromImpute on the other measures (Fig. 4, S8). As a tree-based model, ChromImpute can learn non-linear relationships in the data that PREDICTD cannot, and it is possible that this accounts for some of the difference in performance between the two approaches. Nevertheless, we believe that PREDICTD has additional strengths, including its simpler training procedure, the way the model naturally maps to the problem of imputing the epigenome, and the greater interpretability of the model parameters. In the future we are interested in exploring non-linear models that can implement tensor factorization, particularly deep neural networks. Such models could potentially combine the best of both approaches and lead to even higher quality imputation.

Imputed data represents an important tool for guiding epigenomics studies. It is far cheaper to produce than observed data, closely matches the data observed from experimental assays, and is useful in a number of contexts to generate hypotheses that can be explored in the wet lab. We showed that imputed data can provide insights into ncHARs, and furthermore, Ernst and Kellis [7] previously showed that imputed data tend to have a higher signal-to-noise ratio than observed data, that imputed data can be used to generate high-quality automated genome annotations, and that regions of the genome with high imputed signal tend to be enriched in single nucleotide polymorphisms identified as being enriched in genome-wide association studies (GWAS). Thus, imputed data can provide insight into cell type-specific patterns of chromatin state and act as a powerful hypothesis generator. With just one or two epigenomics maps from a new cell type, PREDICTD can leverage the entire corpus of Roadmap Epigenomics data to generate high qualitypredictions of all assays.

## 4 Methods

### 4.1 Data

We downloaded the consolidated genome-wide signal ( – log_10_ p) coverage tracks in bigWig format from the Roadmap Epigenomics data portal (http://egg2.wustl.edu/roadmap/web_portal/processed_data.html#ChipSeq_DNaseSeq) [2]. These tracks are uniformly processed and currently represent the best-curated collection of epigenomic maps available. In addition, these are the same tracks that Ernst and Kellis [7] used to train ChromImpute, making it easier to compare our modeling approaches.

All observed signal tracks show a higher variance at regions of high signal than at regions of low signal. In order to stabilize this variance across the genome and to make the data more tractable for PREDICTD’s Gaussian error model, we applied an inverse hyperbolic sine transform. This transformation, which has been used in previous studies of epigenomic maps [32], is similar to a log transform but is defined for zero values.

After variance stabilization, we defined five training and test splits such that each data set was in one test set. First, we removed any cell types or assays with fewer than five completed experiments to ensure that there would be enough support for training in each dimension in our model. This left 127 cell types and 24 assays, and a total of 1014 completed experiments (66.6% missing). Next, we split these experiments into five test sets by randomly generating five disjoint subsets of experiments that each contained a stratified sample of experiments from across the available cell types and assays. Thus, in each split 20% of experiments comprise the test set and 80% the training set. In addition to the held out test set, PREDICTD uses additional held out data as a validation set to detect model convergence. To ensure that all data in the training data set contributed equally to the final imputation, the training data for each test fold were further split into eight validation folds by cell type/assay pair so that for any pair of test and validation folds the data split is 20% test (203), 10% validation (100), and 70% training (711). The first four test set folds were used for training and tuning the model parameters, while the last test set fold was held out as a final test set.

Last, the data for each experiment was averaged into 25 bp bins across the genome using the `bedtools map` command [33], and the bins overlapping the ENCODE Pilot Regions and 1kb windows centered at non-coding human accelerated regions were extracted for training the PREDICTD model. The resulting data set contains just over 1.3 million bins, or about 1% of the genome. All experiments reported here were conducted using models trained on this subset of the genome. We find that this is more than enough data to train the model, and imputing the entire genome is a simple matter of applying the learned cell type and assay factors across all positions in the genome.

### 4.2 The SVD3D tensor factorization model

#### 4.2.1 Notation

In the following sections we present the PREDICTD model. As mentioned above, the data set can be represented as a 3D tensor with the axes being the cell types, the assays, and the locations across the genome. We refer to these axes as the cell type, assay, and genome dimensions, respectively. We use capital letters, *J, K*, and *I* to refer to the cardinality of each of these dimensions, and lowercase *j, k, i*, to refer to specific indices in each corresponding dimension. We use the same convention to refer to the number of latent factors in the model, *L*, and individual latent factor indices, *l*. Each dimension has two learned data structures associated with it: a factor matrix, and a bias vector. We use bold capital letters to refer to the factor matrices, and bold lowercase letters to refer to the bias vectors. The cell type factor matrix and bias vector and their dimensions are **C***_J×L_*, and **c**_J×1_, respectively. Similarly, for the assay factor matrix and bias vector: **A***_K×L_*, and a*_K×1_*, and for the genome dimension: **G***_I×L_*, and g*_I×1_*.

#### 4.3 **Model**

In order to perform imputation, we train the PREDICTD tensor decomposition model using the PARAFAC/CANDECOMP procedure [8, 9]. Briefly, in this procedure the three-dimensional tensor is factored into three low-rank matrices, each with the same (user-specified) number of column vectors. These column vectors are called “latent factors,” and the tensor is reconstructed by summing the outer products of the corresponding latent factor vector triplets. More precisely, we start with a three-dimensional data tensor 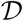 with dimensions *J×K×I*, where *J* = 127 is the number of cell types, *K* = 24 is the number of assays, and *I* = 1,309, 125 is the number of genomic locations (in our case the ENCODE Pilot Regions and 2640 ncHARs at 25 bp resolution), represented by the tensor. This tensor has missing data in fibers along the genome dimension, corresponding to experiments on cell type/assay pairs that have yet to be completed. The completed experiments, corresponding to tensor fibers that contain data, are split into training, validation, and test subsets, or 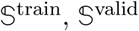, and 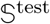, respectively. We factor the tensor 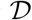, into three factor matrices, and three bias vector terms a, c, and g, that minimize the following objective function:

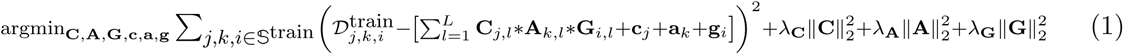

We optimize this objective using parallel stochastic gradient descent [11] with the Adam optimizer [34], and incorporating Nesterov Accelerated Gradient into the optimization as well [35] (Fig. 1).

There are many tensor decomposition methods (reviewed in [27]), however we chose the PARAFAC model because of its relative simplicity. It is not only straightforward to implement and parallelize, but it also requires fewer parameters than other tensor factorization methods [26, 27]. The PARAFAC model also has the nice property that as long as mild conditions hold it will find a solution that is unique in the sense that it is rotationally invariant [26, 27]; this is not a property of other tensor factorization approaches, including Tucker decomposition, which was used in [29].

#### 4.4 Implementation

PREDICTD is implemented in Python 2.7 and built using the Apache Spark 1.6.2 distributed computation framework (http://spark.apache.org). The code is open-source and available on GitHub (https://github.com/tdurham86/PREDICTD.git), and the environment we used to train the model is available on Amazon Web Services as an Amazon Machine Image (see the GitHub repository for info). Models were trained using Amazon Web Services (AWS) Elastic Compute Cloud (EC2) (http://aws.amazon.com) and Microsoft Azure Spark on HDInsight (http://azure.microsoft.com). We bootstrapped an EC2 cluster running Apache Spark 1.6.2 by running the spark-ec2 script (https://github.com/amplab/spark-ec2) on a small EC2 instance (e.g. m3.medium) that we subsequently terminated after the cluster was up and running. Standard cluster configuration was a single m4.xlarge head node instance and one x1.16xlarge worker instance, giving a total cluster size of 2 nodes, 68 cores, and 972 GB of memory. Whenever possible, we used SPOT instances to make the computation more affordable. Microsoft Azure HDInsight clusters had similar resources. All data input to the model and all model output was written to cloud storage; either Simple Storage Service (S3) on AWS, or Blob Storage on Azure.

The data tensor is assembled into a Spark Resilient Distributed Data Structure (RDD) and partitioned among the cluster nodes such that each partition is stored on a single node and contains the data for 1000 genomic loci. This results in about 1300 partitions. The data in each of the 1000 elements in each partition is represented as a `scipy.sparse.csr_matrix` [36] object storing all observed data values for a particular genomic position. Each element of the data RDD also contains the corresponding entries from the **G** factor matrix, g bias vector, and data structures for the Adam optimizer that are specific to each genomic locus (Fig. 1).

The first step of training selects a random 1% of available genomic positions (∼100,000 positions, or ∼0.01% of all 25 bp bins in the genome) for training the cell type and assay parameters. Based on data saturation analysis (Fig. S2), this is enough data to faithfully represent the distribution of data across the genome. The main training phase then proceeds through a series of parallel stochastic gradient descent [11] iterations (Fig. 1C) on this subset of positions. Briefly, at the start of each parallel iteration, copies of the cell type and assay parameters, **C**, **c**, **A**, and a, are sent out to each partition. Each partition undergoes local stochastic gradient descent for 5000 iterations and applies the updates to the local copies of the assay and cell type parameters. The updated cell type and assay parameter values are then passed back to the master node where they are averaged element-wise with the results from all other partitions. The resulting averaged parameters are then copied and distributed to the partitions for the next round of parallel SGD. Note that over all rounds of SGD, we use a learning rate decay schedule of *η_t_* = *η* × (*ϕ_η_*)^*t*–1^, where the learning rate decay parameter *ϕ_η_* = 1 – 1*e*^−6^, and similarly for the Adam first moment parameter: *β1_t_* = *β1 ×* (*ϕ*_β1_)^t–1^, where *ϕ*_*β*1_ = 1 – 1*e*^−6^.

Averaging the parameters after the parallel SGD updates allows the model to share information across the genome dimension; however, the averaging can initially make it harder for the model to converge. The **C** and **A** matrices are initialized randomly from a uniform distribution on the domain (–0.33, 0.33), and thus during the first round of parallel SGD the independent nature of the local updates can lead to inconsistent updates to the latent factors in different partitions. When the results of these inconsistent updates are averaged, they produce poor parameter values, and it then takes many parallel iterations before the parameter values begin to converge. To combat this effect, the main training phase begins with a burn-in stage before attempting parallel SGD. In the burn-in stage, local SGD is performed for one epoch on 8000 genomic loci in a single partition, and after this, the updated **C**, **c**, **A**, and **a** parameters are used in a round of local SGD across the entire 1% training subset to bring the genome dimension up to the same number of updates. This burn-in procedure allows the latent factors to have a consistent initial “identity” across the cluster when starting the parallel SGD updates.

In the parallel SGD stage, the model is trained using the parallel stochastic gradient descent procedure as described above, with 5000 stochastic updates per partition followed by averaging of the updated assay and cell type parameters. Every three parallel SGD iterations, the mean squared error (MSE) is computed for each subset of data (training, validation, and test) and recorded. If the validation MSE is the lowest yet encountered by the model, the parameters from that iteration are copied and saved. Once a minimum number of parallel iterations have completed, the model tests for convergence by collecting the MSE on the validation set for iterations *t –* 35 to *t –* 20 (window 1), and *t* – 15 to *t* (window 2), and using a Wilcoxon rank sum test to determine if window 2 + 1*e*^−5^ > window 1, with one-tailed *p* < 0.05. If this convergence criterion is met, then one of two things happens. First, the model will check whether or not the user has requested a line search on the learning rate. If so, then it will reset the cell type, assay, and genome parameters to those found at the iteration with the minimum validation MSE and resume parallel SGD after halving the learning rate and reducing the Adam first moment weight *β*1_new_ = *β*1_old_ – (1.0 – *β*1_old_). When training the model, we used a line search of length three, so the model was restarted from the current minimum and learning rate halved and *β*1 adjusted three times. Once the line search is complete, or if no line search was requested, then the model stops parallel SGD, fixes the assay and cell type parameters, and finishes training on the genome parameters only.

Once the main phase of training is complete, the last phase of model training applies the cell type and assay parameters across all genomic positions. This is accomplished by fixing the cell type and assay parameters and calculating the second order solution on the genome parameters only. This requires a single least squares update per genomic position, and it is possible because fixing the cell type and assay parameters makes our objective function convex over the genome parameters. Once the final genome parameters are calculated, the assay, cell type, and genome parameters are saved to cloud storage, and the imputed tensor is computed and saved to the cloud for further analysis. On average, the entire training takes about 750 parallel iterations, and about 6 hours (wall clock time).

The above procedure is executed for every validation fold in a given test fold, and then the final imputed values for the held-out test data sets are calculated as a simple average of the corresponding imputed values from each validation fold. Thus, for the results we report here, each imputed value represents the consensus of eight trained models.

**Table 2:** Hyperparameter values. The third column indicates whether the hyperparameter value was selected using Spearmint, and an asterisk indicates the final value was tuned by hand after Spearmint optimization.

| Hyperparameter | Value | Spearmint? |
| --- | --- | --- |
| $\eta$ | 0.005 | Y |
| $\phi_\eta$ | $1 - 1e^{-6}$ | N |
| $\beta_1$ | 0.9 | N |
| $\phi_{\beta_1}$ | $1 - 1e^{-6}$ | N |
| $\beta_2$ | 0.999 | N |
| $L$ | 100 | Y* |
| $\lambda_{\mathbf{C}}$ | 3.66 | Y |
| $\lambda_{\mathbf{A}}$ | $1.23e^{-7}$ | Y |
| $\lambda_{\mathbf{G}}$ | $1.23e^{-5}$ | Y |
| $\lambda_{\mathbf{G}_2}$ | 2.9 | N |

##### 4.4.1 Hyperparameter Selection

One of the challenges of working with this type of model is that there are many hyperparameters to tune. These include the number of latent factors *L*, the learning rate *η*, the learning rate decay *ϕ_η_*, the Adam first moment coefficient *β*_1_ and its decay rate *ϕβ*_1_, and a regularization coefficient for each latent parameter matrix (λ**_A_**, λ**_C_**, λ**_G_**). This hyperparameter space is very large, so instead of implementing a simple grid search, we used a software package called Spearmint [37] to dynamically optimize the hyperparameters by fitting a Gaussian process model to the hyperparameter space and finding the minimum. After Spearmint picked a combination of parameters that gave a good solution, then we fine-tuned them by hand. Our final hyperparameter values for model training are listed in Table 2.

##### 4.4.2 Imputing the whole genome

Although for the results reported here we only imputed about 1% of the genome, we are in the process of extending our imputed tracks to the whole genome. This simply involves applying the cell type and assay parameters across the whole genome in a procedure identical to the final stage of training the model. We will make these imputed data sets available through ENCODE.

##### 4.4.3 Imputing data for a novel cell type

Given data from a modeled assay from a cell type that the model has not seen before, the factor matrices must be re-trained. We showed that such a new cell type can be accurately imputed with observed data for just a single assay (Fig. 2C) when we simulated such a case by training the model on all of the observed data except for some tracks for the “CD3 Primary Cells Cord Blood” cell type. We trained a model in which H3K4me1 was the only assay from this cell type included in the training set, as well as experiments with only H3K4me3 included, and with H3K4me3 and H3K9me3 included. Model training proceeded as described above, except we used only four validation folds in each of these cases to expedite model training.

##### 4.4.4 Advantages of the consumer cloud

Cloud computing is becoming a powerful tool for bioinformatics. Large consortia such as the Encyclopedia of DNA Elements [1] and The Cancer Genome Atlas (http://cancergenome.nih.gov) are making their data available on cloud platforms. As computational analyses grow more complex and require more computing resources to handle larger data sets, the cloud offers two distinct advantages. First, cloud services provide a centralized way to host large data sets used by the community that makes data storage, versioning, and access more simple and efficient. Transferring gigabytes, or even terabytes, of data is slow and expensive in terms of network bandwidth, but moving code and computation to the data is fast and cheap. Second, in addition to hosting data sets, cloud services can host saved computing environments. Such virtual machine images can help with reproducibility of results for complex analyses because the code can be written in such a way that other users can not only use the same code and data as the original authors, but they can run the analysis in the same computing environment. One downside of cloud computing for labs that have access to a local cluster is that cloud resources are charged by usage; nevertheless, generating high quality imputed data using PREDICTD is extremely cost effective compared to collecting the observed data. Imputing the data for this paper cost on the order of US $3 per data set, which is more than two orders of magnitude lower than the cost of completing these experiments in the wet lab, and this cost can be expected to drop as computational resources become cheaper and more efficient optimization methods are devised.

#### 4.5 Imputation quality metrics

##### 4.5.1 Visual inspection

We generated tracks for the imputed data by extracting the data for each 25 bp bin from the imputed results, writing the results to file in bedGraph format, then converting to bigWig using the bedGraphToBigWig utility from UCSC. Imputed tracks were visually inspected alongside Roadmap Consolidated data tracks and peak calls in the UCSC Genome Browser. We did not reverse the variance stabilizing inverse hyperbolic sine transform when evaluating model performance. This is appropriate because it maintains the Gaussian error model that underlies the PREDICTD optimization.

##### 4.5.2 Quality metrics

We implemented ten different quality assessment metrics (listed below), the last seven of which were first reported for ChromImpute [7]. We report these metrics computed over the ENCODE Pilot Regions S8.

- Mean squared error over all available genomic positions (**MSEglobal**)
- Mean squared error over the top 1% of windows based on observed signal (**MSE1obs**)
- Mean squared error over the top 1% of windows based on imputed signal (**MSE1imp**)

–**MSE1imppred:** Mean squared error over the top 1% of windows based on PREDICTD signal
–**MSE1impchrimp:** Mean squared error over the top 1% of windows based on chromImpute signal
–**MSE1impme:** Mean squared error over the top 1% of windows based on main effects signal
- Pearson correlation over all available genomic positions (**GWcorr**)
- Percentage of top 1% of observed signal windows that are also in the top 1% of imputed windows (**Match1**)
- Percentage of top 1% of observed windows found in the top 5% of imputed windows (**Catch1obs**)
- Percentage of top 1% of imputed windows found in the top 5% of observed windows (**Catch1imp**)
- Recovery of the top 1% of observed windows from all imputed windows based on the area under the curve of the receiver operating characteristic (**AucObs1**)
- Recovery of the top 1% of imputed windows from all observed windows based on the area under the curve of the receiver operating characteristic (**AucImp1**)

#### 4.6 Analyzing model parameters

The parameter values corresponding to individual latent factors are not individually interpretable, but intuitively we can understand that each latent factor describes some pattern in the data that the model finds useful for imputation. For example, the first latent factor (i.e., column 0 in each of the three factor matrices) might contain values that capture a pattern of high signal in promoter marks, in blood cell types, at active genes. In such a case the value at this latent factor for a particular assay might suggest how often that mark is found at promoters; for a particular cell type its relatedness to blood; and for a genomic locus how many promoter-associated features occur there in blood cell types. If these three conditions hold, then the model is likely to have more extreme values for these parameters that end up imputing a high value for that cell type/assay pair at that genomic position.

##### 4.6.1 Clustering cell types and assays

The rows of the cell type and assay factor matrices, with each row containing the model parameters for a particular cell type or assay, respectively, were clustered using hierarchical clustering. This analysis was implemented in Python 2.7 using `scipy.spatial.distance.pdist with metric=’cosine’` to generate the distance matrix, and `scipy.cluster.hierarchy.linkage with method=’average’` to generate clusters. The columns of each factor matrix (i.e. the latent factor vectors) were also clustered in the same way to help with visualizing the clusters. The parameter values were plotted as a heat map with rows and columns ordered according to the results of the hierarchical clustering.

##### 4.6.2 Finding signatures of genomic elements

The genome factor matrix is too large to usefully visualize as a heatmap, so we sought to find patterns correlating with known genomic features to summarize the parameters. We mapped all annotated protein-coding genes from the GENCODE v19 human genome annotation [12] (https://www.gencodegenes.org/releases/19.html) with a designated primary transcript isoform (called by the APPRIS pipeline) to a canonical gene model consisting of nine components: promoter, 5’ UTR, first exon, first intron, middle exon, middle intron, last exon, last intron, and 3’ UTR. The promoter for each gene was defined as the 2 kb region flanking the 5’ end of the gene annotation, while the other components were either taken directly from the GENCODE annotation (5’ UTR, exons, 3’ UTR) or were inferred (introns). For each gene, each component was split into ten evenly spaced bins and the signal for each latent factor was averaged so that there was a single value for each latent factor for each bin. Coding regions for genes with a single exon or two exons were mapped only to first exon, or first exon and last exon components, respectively. Genes with only one or two introns were handled analogously. For genes with multiple middle exons and introns, each exon/intron was binned independently and the data for each middle exon/intron bin was averaged across all middle exons/introns. In order to plot the results, outlier values in the bins (defined as any values outside 1.5 * IQR) were removed and the remaining values averaged across corresponding bins for all binned gene models. This resulted in a matrix containing latent factors on the rows and gene model bins on the columns. The latent factors (rows) were clustered using hierarchical clustering, with `scipy.spatial.distance.pdist(metric=’euclidean’)` to generate the distance matrix and `scipy.cluster.hierarchy.linkage(method=’ward’)` to generate clusters, and this matrix was plotted as a heat map.

To compile a reference list of genome coordinates containing distal regulatory elements that is orthogonal to our imputed data, we downloaded P300 peak data from six ENCODE cell lines (A549, GM12878, H1, HeLa, HepG2, and K562), filtered for peaks with FDR < 0.01, merged the peak files with `bedtools merge` to create a single reference list, and averaged genome latent factor values as in the gene model explained above for ten 200 bp bins covering 2kb windows centered on these peaks.

To validate that the patterns in the genome parameters were not due to chance, we generated the same heatmap, but before averaging the bins for each gene model and P300 site we randomly permuted the order of the genome latent factors (Fig. S6). All signal disappears except for a band of lower parameter values at the P300 peak centers, which implies that the bins closer to the peak centers have a bias toward lower values across most of the latent factors.

#### 4.7 Comparing to ChromImpute

To compare the performance of PREDICTD with ChromImpute, we downloaded the ChromImpute results from the Roadmap Epigenomics website and put them through the same pipeline as for the observed data: Convert to bedgraph, use `bedtools map` to calculate the mean signal over 25 bp bins, extract the bins overlapping the ENCODE Pilot Regions, apply the inverse hyperbolic sine transform, and store the tracks in a Spark Resilient Distributed Dataset (RDD) containing a list of `scipy.sparse.csr_matrix objects`.

We calculated all of the quality metrics on these ChromImpute data sets and plotted these results against those for PREDICTD for each experiment as a ternary scatter plot (Fig. 4, S8). We also averaged each element of this ChromImpute RDD with its corresponding element in the PREDICTD results, calculated the quality metrics, and compared against ChromImpute alone (Fig. 4). In order to compare both ChromImpute and PREDICTD to the baseline main effects model, we used ternary plots [38] to project the three dimensional comparison of each experiment to two dimensions. Each point on these ternary plots represent the relative magnitude of each dimension for that point. So, each coordinate (*x,y,z*) in Cartesian space is projected to a point (*x’, y’, z’*) such that 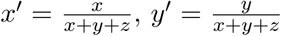, and 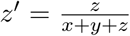. Thus, for the case where *x* = *y* = *z* the corresponding point (x’, y’, z’) = (0.33, 0.33, 0.33) and will fall at the center of the ternary plot, while points that lie along the Cartesian axes will fall at the extreme points of the ternary plot (e.g. (*x*,*y*,*z*) = (1, 0, 0) = (*x*’,*y*’,*z*’).

It is important to emphasize that the quality measure with the best PREDICTD performance, MSEglobal, is also explicitly optimized by the PREDICTD objective function during training. This shows that PREDICTD is doing well on its assigned learning task, and highlights the importance of designing an objective function that reflects the task that the model will address. As such, it should be possible to tune the objective function to perform better on other quality measures if need be. For example, in an attempt to boost PREDICTD’s performance on regions with higher signal we experimented with weighting genomic positions by ranking them by the sum of their signal level ranks in each training data set. This provided some improvement on the MSE at the top 1% of observed signal windows measure (MSE1obs), but we ultimately decided to pursue the simpler and more balanced objective function presented here.

#### 4.8 Assessing enhancer signatures at non-coding human accelerated regions

##### 4.8.1 Clustering ncHARs and cell types

We downloaded the non-coding human accelerated region coordinates used in Capra, 2013 [10], removed any that overlapped a protein-coding exon according to the GENCODE v19 annotations [12], and extracted all available observed and imputed data for the enhancer-associated assays H3K4me1, H3K27ac, and DNase at these regions. Some cell types were lacking observed data for H3K27ac (29) and/or DNase (74), but observed data for H3K4me1 was available in all cell types. We took the mean signal for each assay at each ncHAR coordinate and used that as input to the subsequent analysis.

First, we extracted the first principal component of the three assays for all ncHARs and cell types using `sklearn.decomposition.TruncatedSVD` [39] to reduce the assay dimension length from three to one and construct a matrix of ncHARs by cell types. This also had the effect of filling in missing values for the observed data. Next we wanted to cluster the ncHARs and cell types, and so we first used the matrix based on imputed data to assess how many clusters would be appropriate for the data. Briefly, for both the ncHAR and cell type dimensions, we conducted an elbow analysis by calculating the Bayesian information criterion (BIC) for k-means clustering results for all values 2 <= *k* <= 40, as well as a silhouette analysis on the same range of values for k (Fig. S9). Based on the results, we decided that *k* = 6 for both dimensions was a good balance of distance between clusters and number of clusters.

Next, we clustered the imputed and observed matrices with the scikit-learn `sklearn.cluster.bicluster.SpectralBiclustering` class [39] to generate a biclustering using six column clusters and six row clusters. And finally, we plotted the clustering results for the imputed data as a heatmap in which each cell is the sum of the mean H3K4me1, H3K27ac, and DNase signal at a particular ncHAR in a particular cell type. We also plotted the tissue assignments for ncHARs with predicted developmental enhancer activity based on EnhancerFinder [19] calls in the Capra, 2013 [10] paper alongside our ncHAR clusters (Fig. 5A). The same plot for the observed data is shown in Fig. S10.

##### 4.8.2 Gene Ontology analysis of cluster results

We found the nearest gene for each ncHAR by using `bedtools closest` with the GENCODE v19 human genome annotation [12]. Then, we grouped these genes by our ncHAR clustering results on imputed data and searched for Gene Ontology biological process term enrichments with the PANTHER [24, 25] web interface (http://pantherdb.org) using the overrepresentation test (release 20160715) with default parameters (specific details: Annotation Version and Release Date - PANTHER 11.1 Released 2016-10-24; Reference List - Homo sapiens (all genes in database); Annotation Data Set - PANTHER GO-Slim Biological Process; we used Bonferroni correction for multiple testing). Four of our clusters returned terms with Bonferroni-corrected *p* < 1.0, and we report all such terms here (Fig. 5B, S11).

##### 4.8.3 Assessing clustering performance

In order to compare the clustering results on the imputed data to the observed data, we used the adjusted Rand index, which assesses how often pairs of data points are put in the same or different clusters, on the ncHAR and cell type clusters independently. we also conducted the same clustering analysis on the observed data after shuffling the ncHARs to other non-coding coordinates (Shuffled), and after switching out the enhancer-associated marks for H3K4me3, H3K36me3, and H3K27me3, which are not associated with enhancers, and compared the resulting clusters with the enhancer-associated imputed data and observed data clusters, again using the adjusted Rand index. Last, we used the adjusted Rand index once more to assess the similarity of our biclustering results to the grouping of ncHARs based on predicted tissue-specific developmental enhancer activity from Capra, 2013 [10] (Fig. 5B).

## 5 Acknowledgements

We gratefully acknowledge support from the Amazon Web Services Cloud Credits for Research program and Microsoft Azure for Research program for providing computing cycles to help with the development of PREDICTD. This work was funded by National Institutes of Health awards R01 ES024917 and U41 HG007000.

---

**Figure S1:**
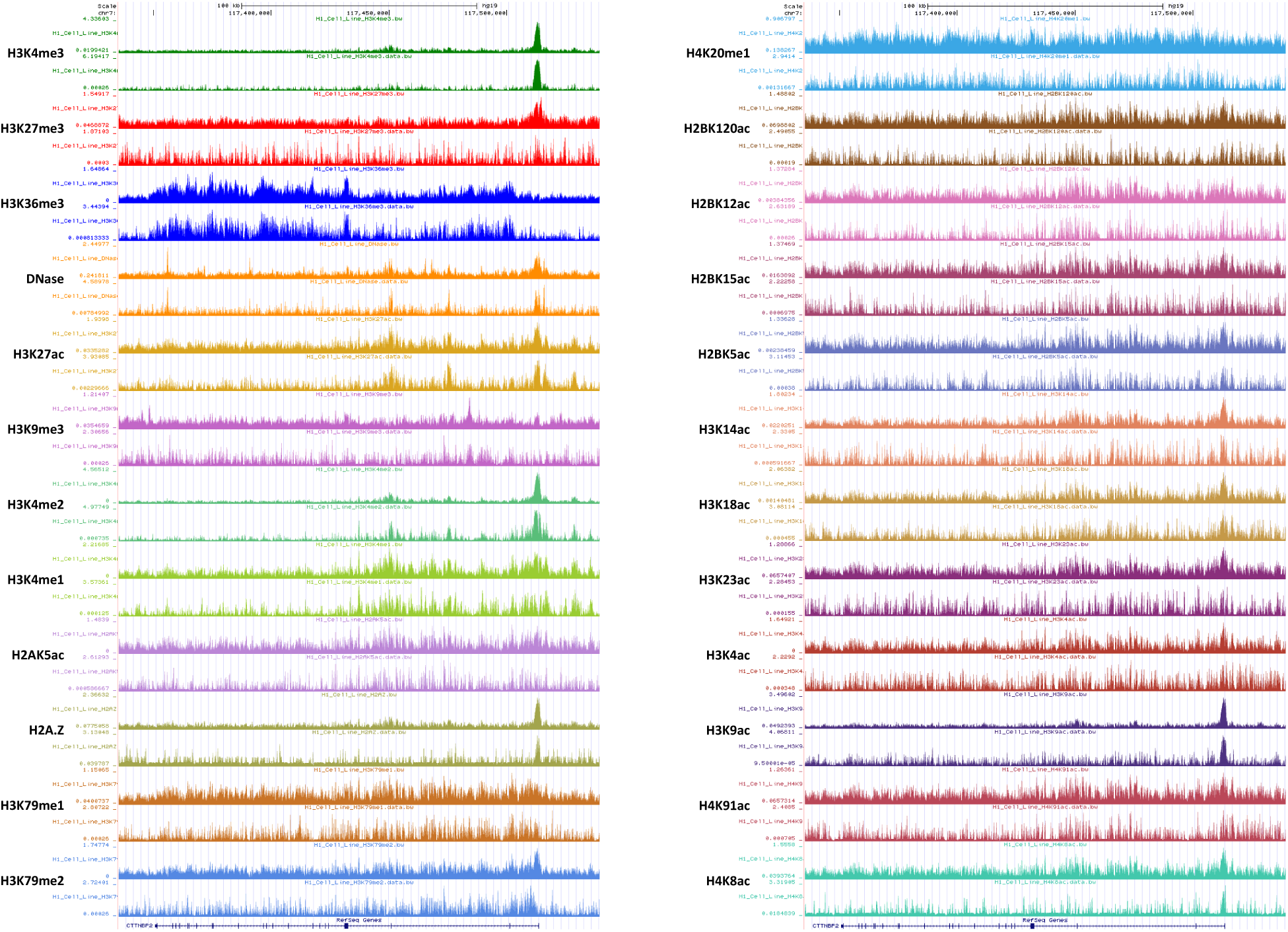
Tracks showing paired PREDICTD (top) and observed (bottom) data for the H1 Cell Line cell type, which is one of only three cell types for which observed data are available for all assays. The signal is variance stabilized with the inverse hyperbolic sine transform, and the tracks are auto-scaled by the genome browser to highlight the shape of the signal.

**Figure S2:**
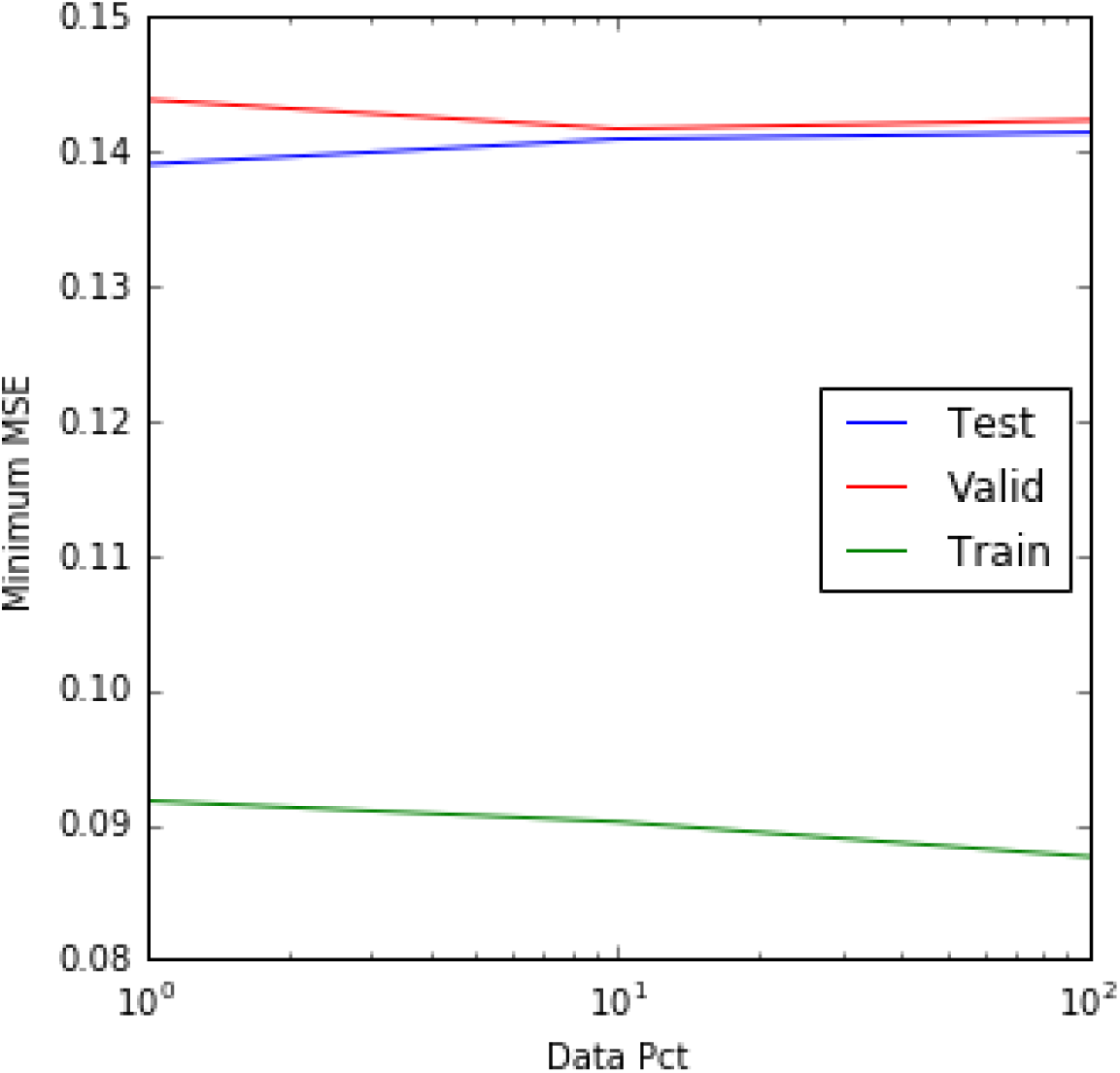
Starting with 1% of the genome (all ENCODE Pilot Regions), data saturation curves on the first training/test split show very stable performance even over a reduction in data of two orders of magnitude (1.0% to 0.01% of the genome). In order to speed training of the cell type and assay parameters during the parallel stochastic gradient descent phase, we trained all models for this paper on 0.01% of the genome (about 100,000 genomic positions).

**Figure S3:**
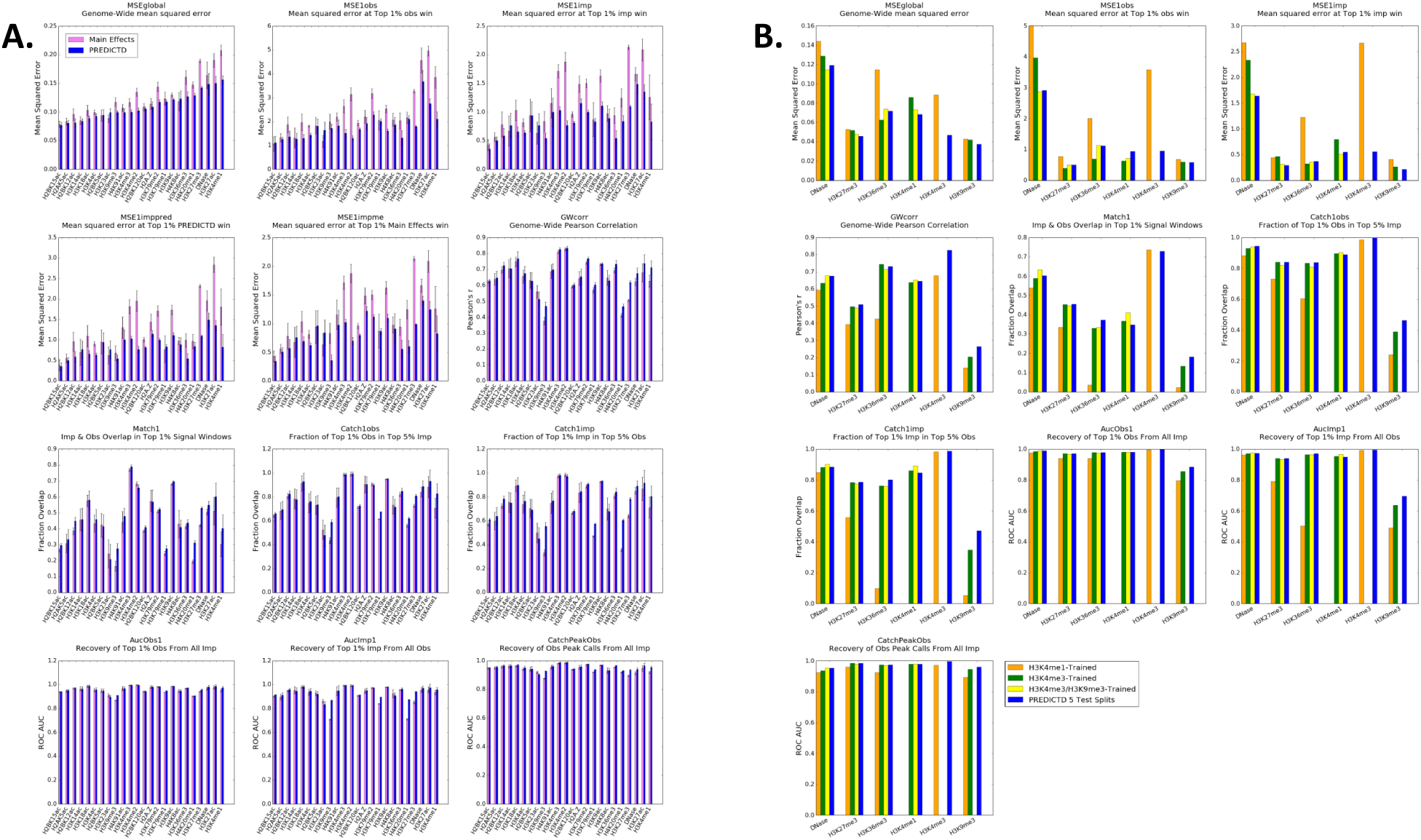
**A.** Plots for all metrics evaluating the performance of the main effects baseline vs PREDICTD. Bars show the mean value for each quality metric over all available observed data sets for each assay; error bars show the standard error of the mean. **B.** Plots for all metrics evaluating the performance of the model on a cell type with only certain assays included in the training set. Each bar indicates the quality measure value for that assay in a single cell type, “CD3 Primary Cells from Cord Blood.

**Figure S4:**
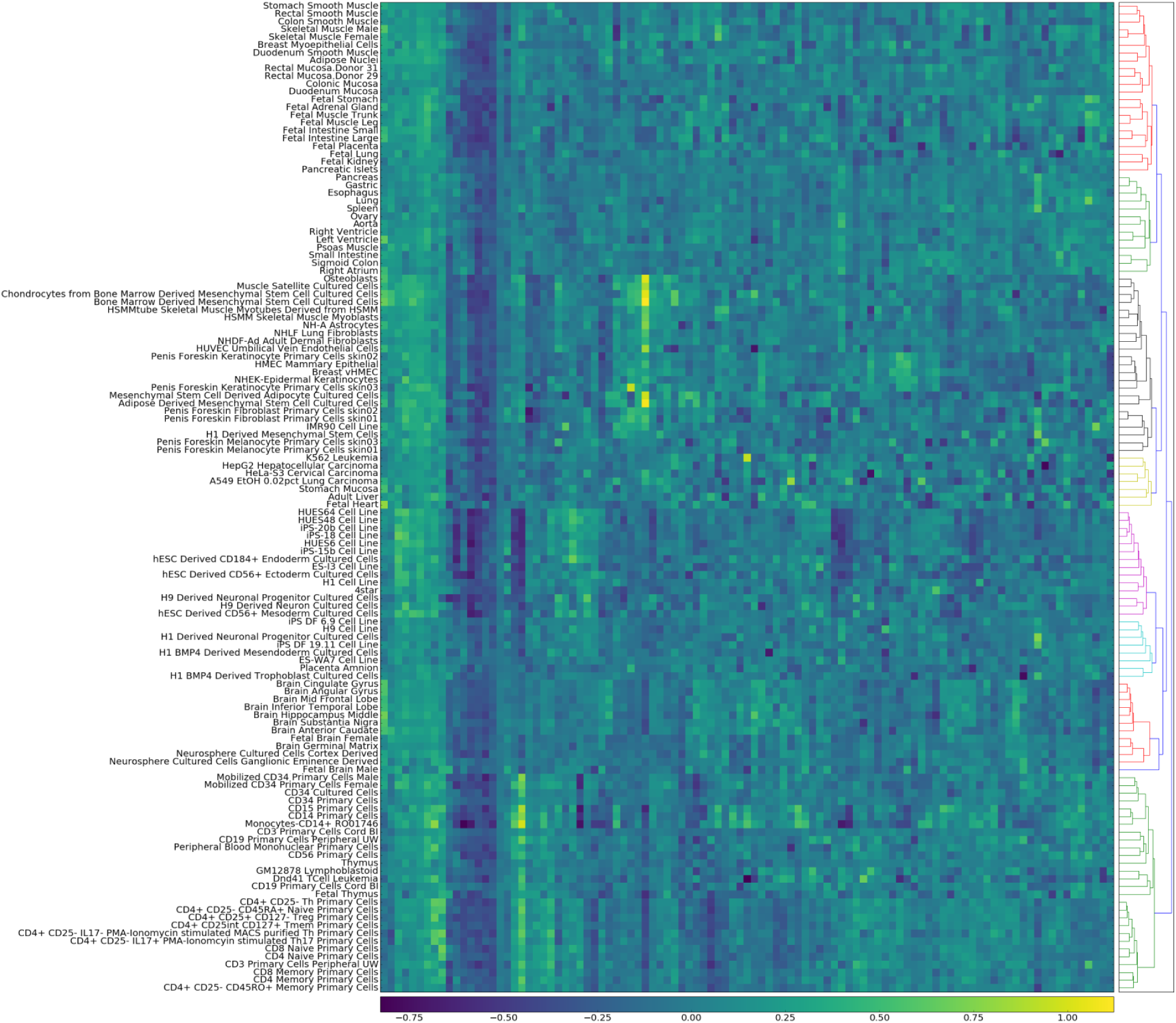
Hierarchical clustering of all cell types based on latent factor parameter values. For legibility, Fig. 3A shows a clustering of only the lower 62 rows of this figure.

**Figure S5:**
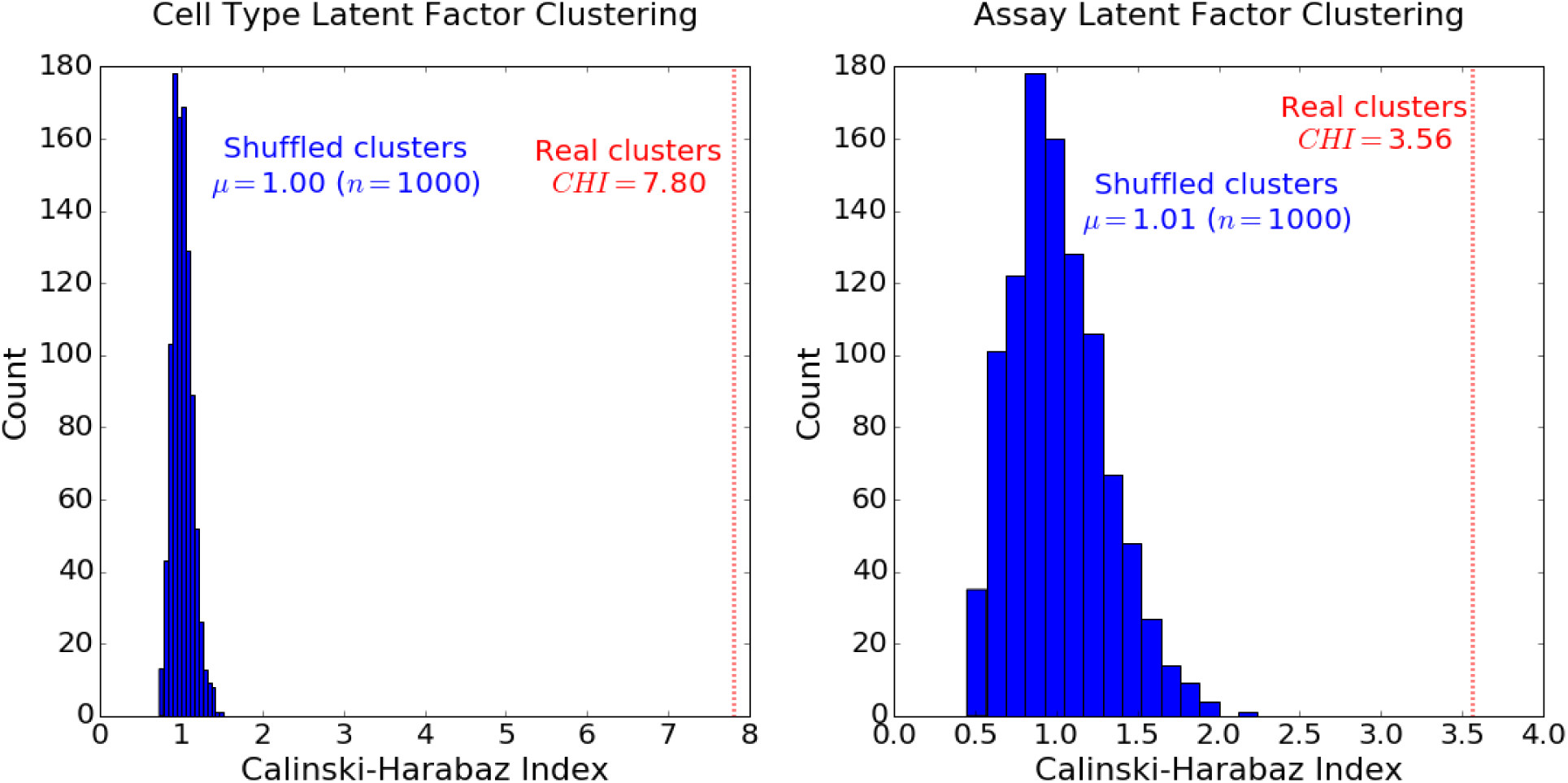
Hierarchical clustering of cell types and assays by latent factor parameter values results in much better cluster separation than randomly assigning cluster identities. A higher value on the Calinski-Harabaz Index indicates that the clusters are denser and better-separated. Linkage trees were cut at eight clusters for cell types and four clusters for assays.

**Figure S6:**
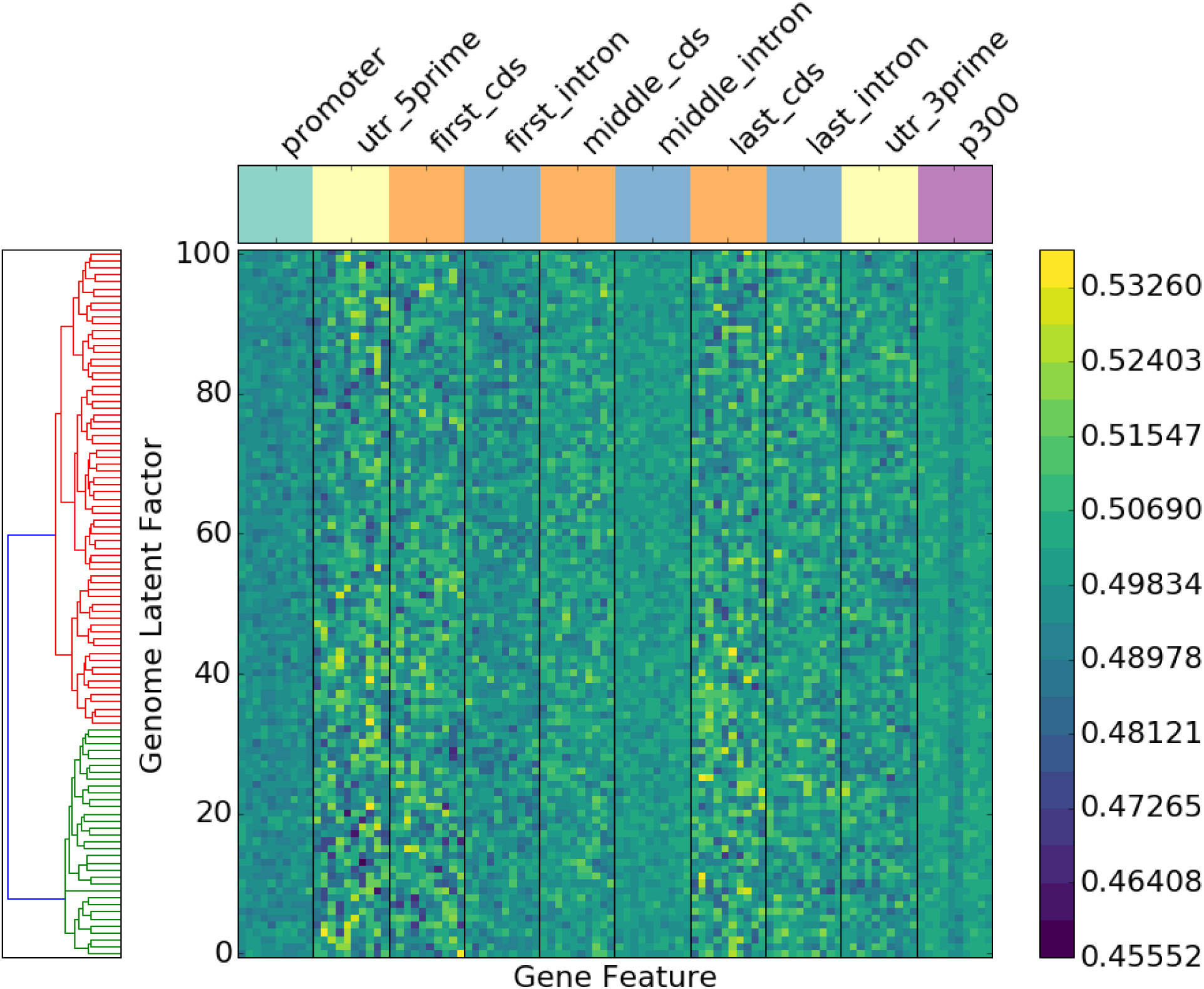
The same analysis was completed as in Fig. 3C, but after randomly permuting the latent factors at each genomic position. Randomly permuted latent factors do not show the same patterns at genomic elements.

**Figure S7:**
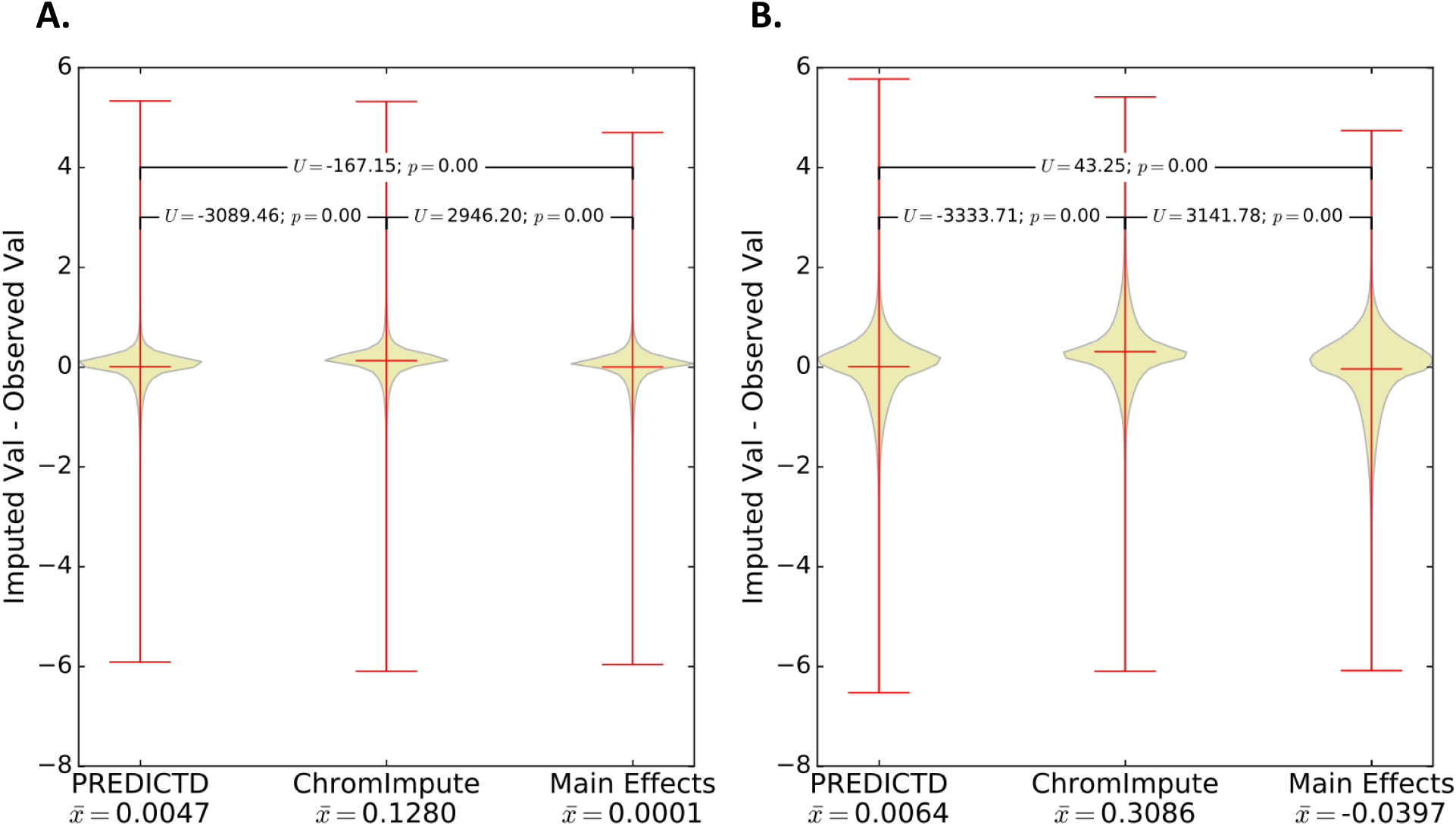
The error distribution of ChromImpute values is more positive than that of PREDICTD. This means that on average ChromImpute tends to over-estimate signal amplitude compared to PREDICTD. Error distributions for each model were compared with the Mann-Whitney U test, and the sample mean is reported for each model on the x-axis. **A.** Random sample of 100,000 genomic positions. **B.** Top 100,000 genomic positions by summing all observed data values at each position.

**Figure S8:**
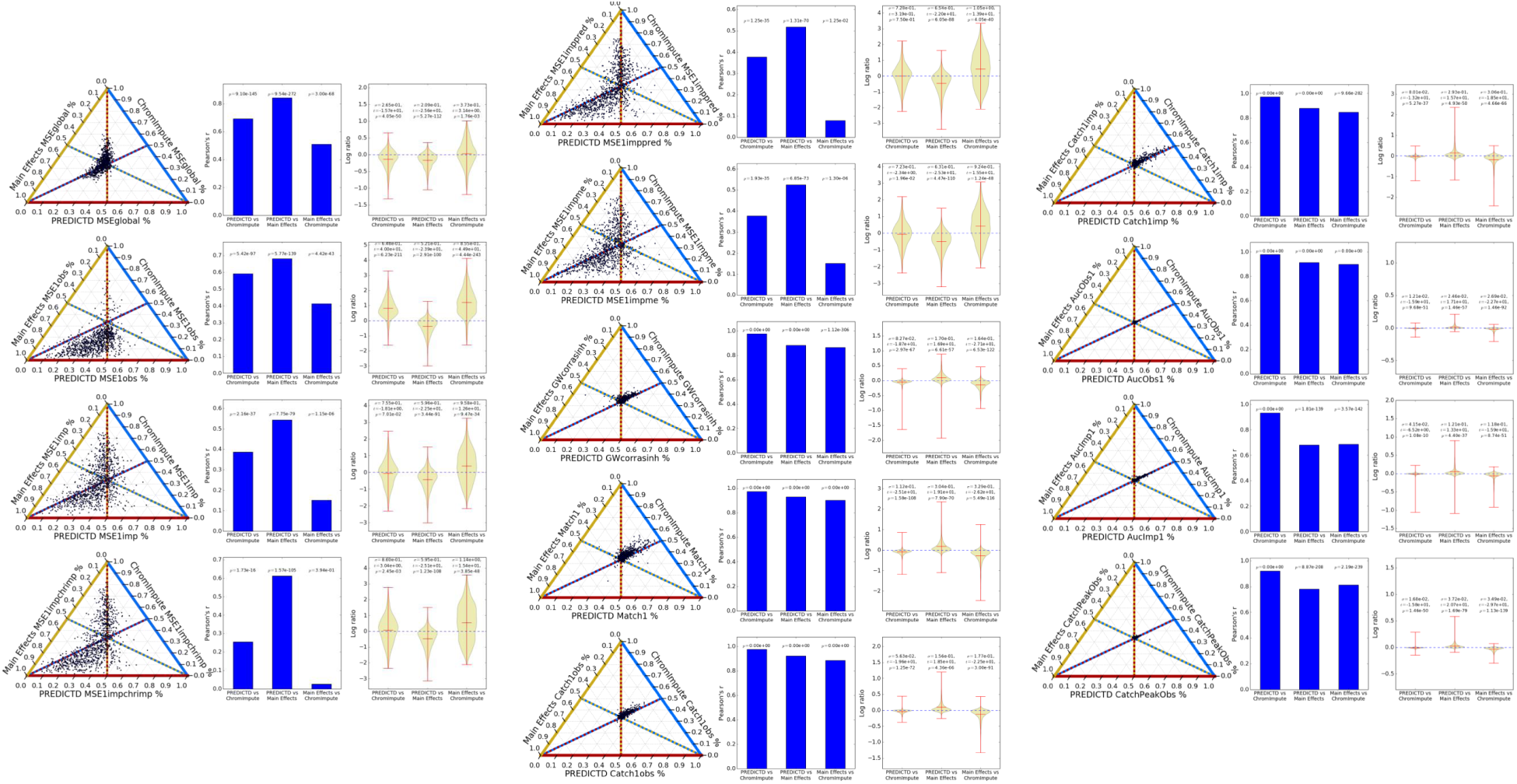
Comparison of all metrics for PREDICTD, ChromImpute, and Main Effects models. The first plot in each triplet is the ternary plot, as described in Fig. 4, the second is the Pearson correlation between the metric values for each pair of models, and the third is the distribution of the natural log fold-change in metric value between corresponding experiments in pairs of models. Note that the correlation of PREDICTD with ChromImpute is always higher than the correlation between Main Effects and ChromImpute, indicating that PREDICTD tends to agree more with ChromImpute than Main Effects does. In addition, the mean log fold change between PREDICTD and ChromImpute is always either closer to zero than the log fold change between Main Effects and ChromImpute, indicating more comparable quality measure values between PREDICTD and ChromImpute, or the mean log fold change indicates stronger performance by PREDICTD (MSEglobal, MSE1imp, and MSE1impme) than ChromImpute. In all, the quality measures show that PREDICTD performs very similarly to ChromImpute, and more so than Main Effects does.

**Figure S9:**
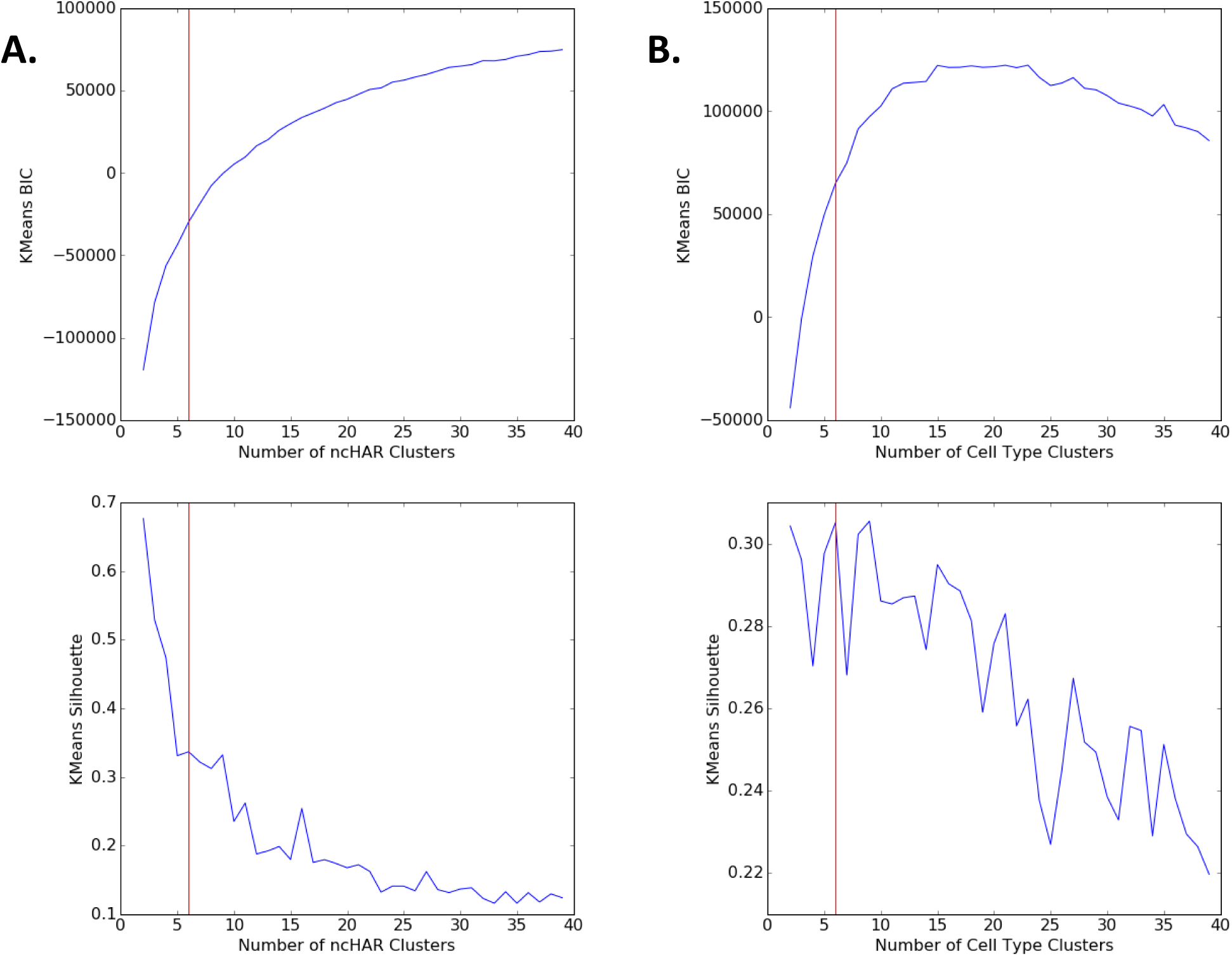
To pick the number of ncHAR and cell type clusters, we used k-means clustering on the rows (ncHARs) and columns (cell types) of the biclustering input matrix (see Methods) for imputed data and conducted a Bayesian Information Criterion (BIC) elbow analysis, as well as a silhouette score analysis. Assessment of the quality of the clustering is based on finding a balance that achieves an appropriate number of clusters that are still well-separated. Elbow analysis based on BIC, as well as silhouette analysis suggests that 6 (vertical line) is a reasonable number of clusters for both ncHARs (**A.**) and cell types (**B.**); this point is at about the maximum of the second derivative in the elbow plots and also near a local maximum in the silhouette plots.

**Figure S10:**
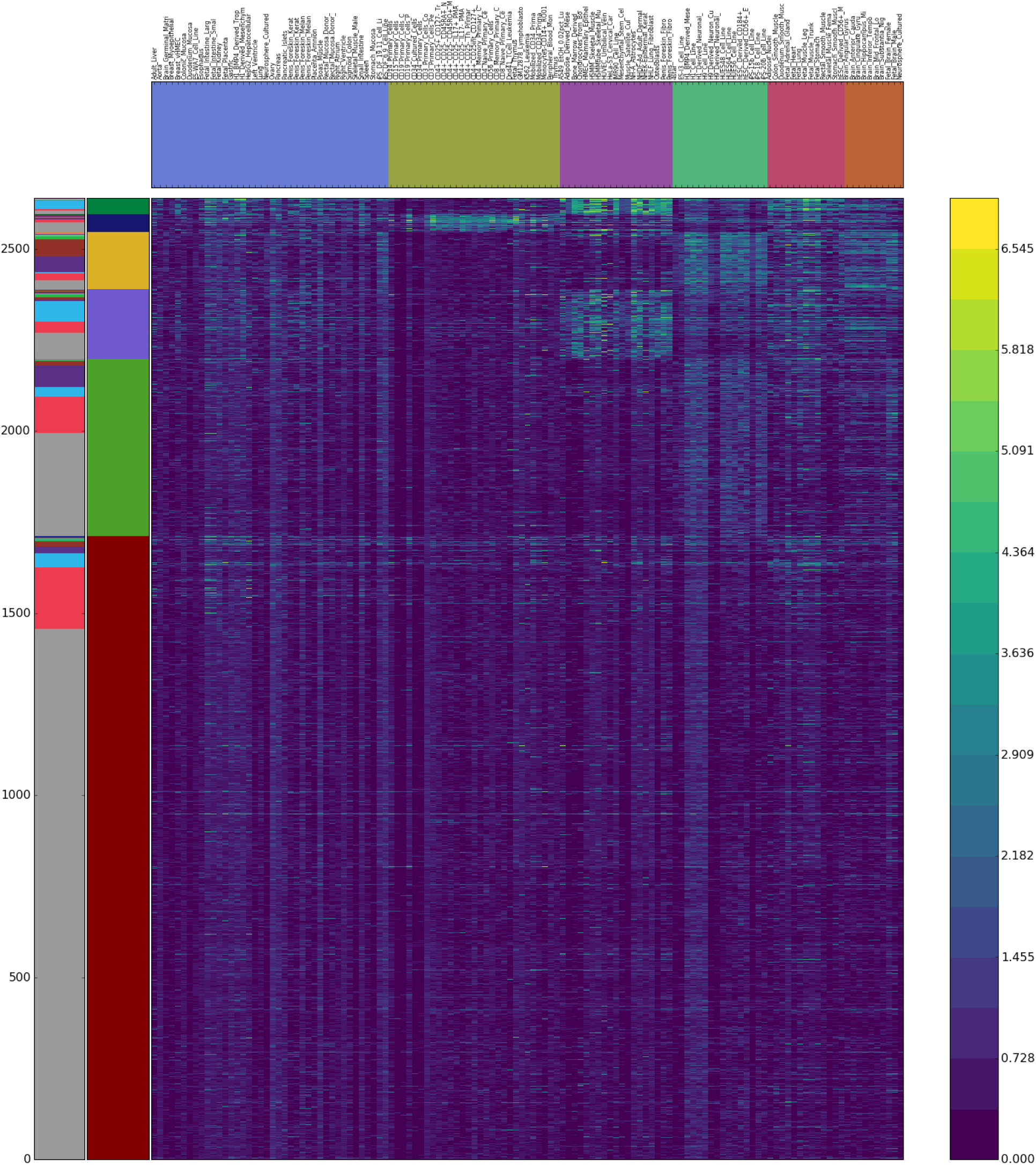
Heatmap showing biclustering results and signal from observed data at ncHARs for the H3K27ac, H3K4me1, and DNase assays. The clustering results are very similar to those for imputed data reported in Fig. 5A.

**Figure S11:**
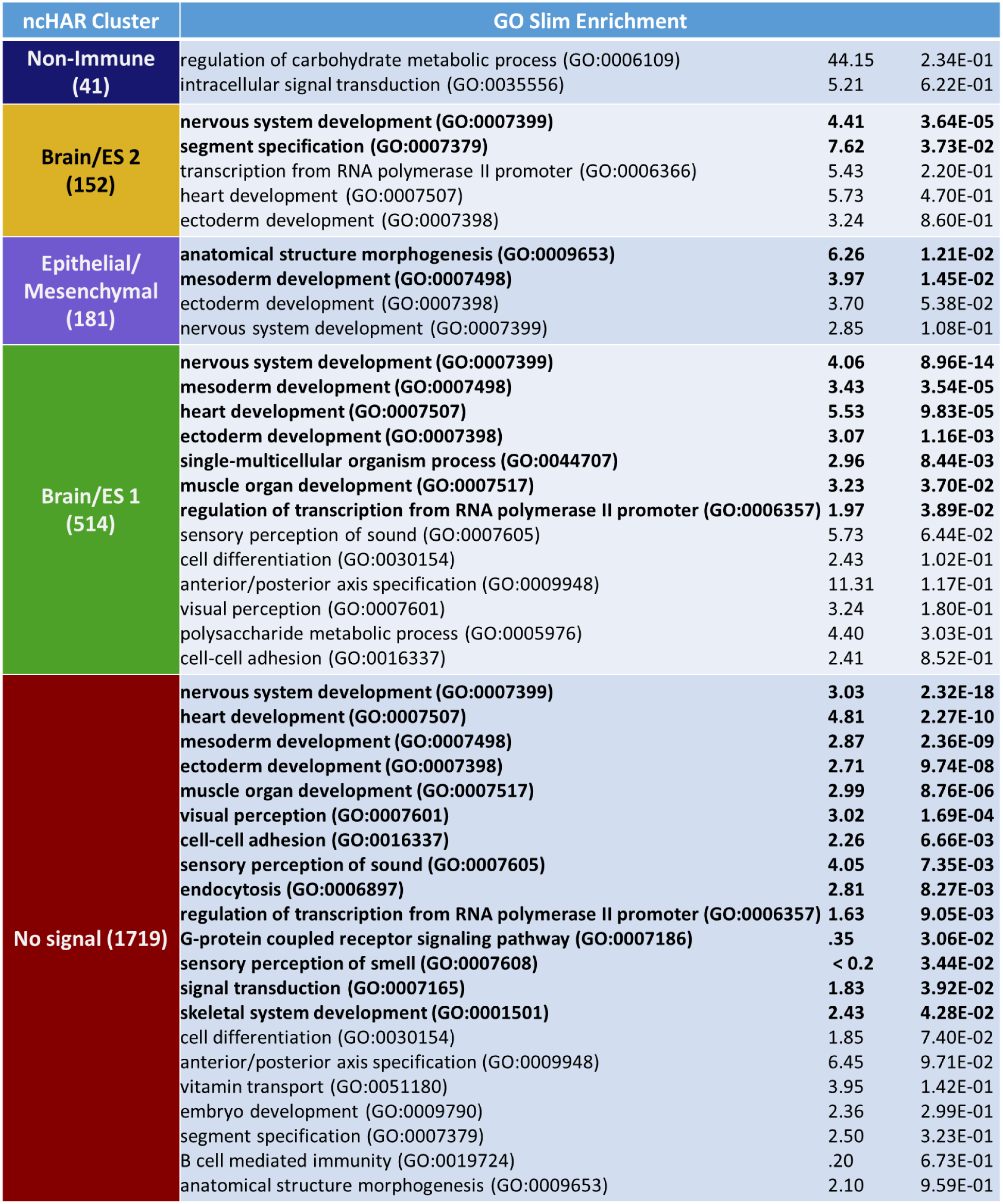
All Gene Ontology terms for the list in Fig. 5B with Bonferroni-corrected p-values less than 1.0. Terms with *p* <0.05 are in bold.

